# Renal L-2-hydroxyglutarate dehydrogenase activity promotes hypoxia tolerance and mitochondrial metabolism in *Drosophila melanogaster*

**DOI:** 10.1101/2024.05.18.594772

**Authors:** Nader H. Mahmoudzadeh, Yasaman Heidarian, Jason P. Tourigny, Alexander J. Fitt, Katherine Beebe, Hongde Li, Arthur Luhur, Kasun Buddika, Liam Mungcal, Anirban Kundu, Robert A. Policastro, Garrett J. Brinkley, Gabriel E. Zentner, Travis Nemkov, Robert Pepin, Geetanjali Chawla, Sunil Sudarshan, Aylin R. Rodan, Angelo D’Alessandro, Jason M. Tennessen

**Affiliations:** Department of Biology, Indiana University, Bloomington, IN 47405, USA; Department of Internal Medicine, Division of Nephrology and Hypertension, and Molecular Medicine Program, University of Utah, Salt Lake City, UT 84112, USA; Medical Service, Veterans Affairs Salt Lake City Health Care System, Salt Lake City, UT, USA; Department of Urology, University of Arizona in Tucson, AZ, USA; Department of Urology, University of Alabama at Birmingham, Birmingham, AL, USA; University of Colorado Denver - Anschutz Medical Campus, Aurora, Colorado, USA; Department of Chemistry, Indiana University, Bloomington, IN 47405, USA; Department of Life Sciences, School of Natural Sciences, Shiv Nadar Institute of Eminence, Dadri, Uttar Pradesh 201314, India; Affiliate Member, Melvin and Bren Simon Cancer Center, Indianapolis, IN, 46202, USA

**Keywords:** L-2-hydroxyglutarate, L2hgdh oncometabolite, hypoxia, *Drosophila*

## Abstract

The mitochondrial enzyme L-2-hydroxyglutarate dehydrogenase (L2HGDH) regulates the abundance of L-2-hydroxyglutarate (L-2HG), a potent signaling metabolite capable of influencing chromatin architecture, mitochondrial metabolism, and cell fate decisions. Loss of L2hgdh activity in humans induces ectopic L-2HG accumulation, resulting in neurodevelopmental defects, altered immune cell function, and enhanced growth of clear cell renal cell carcinomas. To better understand the molecular mechanisms that underlie these disease pathologies, we used the fruit fly *Drosophila melanogaster* to investigate the endogenous functions of L2hgdh. Our studies revealed that while L2hgdh is not essential for growth or viability under standard culture conditions, *L2hgdh* mutants are hypersensitive to hypoxia and expire during the reoxygenation phase with severe disruptions of mitochondrial metabolism. Moreover, we find that the fly renal system (Malpighian tubules; MTs) is a key site of L2hgdh activity, as *L2hgdh* mutants that express a rescuing transgene within the MTs survive hypoxia treatment and exhibit normal levels of mitochondrial metabolites. We also demonstrate that even under normoxic conditions, *L2hgdh* mutant MTs experience significant metabolic stress and are sensitized to aberrant growth upon Egfr activation. Overall, our findings present a model in which renal L2hgdh activity limits systemic L-2HG accumulation, thus indirectly regulating the balance between glycolytic and mitochondrial metabolism, enabling successful recovery from hypoxia exposure, and ensuring renal tissue integrity.

## INTRODUCTION

L-2-hydroxyglutarate (L-2HG) is a potent signaling molecule that acts as a competitive inhibitor of α-ketoglutarate (αKG)-dependent dioxygenases (Chowdhury et al., 2011, Xu et al., 2011, Du and Hu, 2021). As a result, ectopic L-2HG accumulation can interfere with activity of the tricarboxylic acid (TCA) enzyme α-ketoglutarate dehydrogenase, the Jmj-class of histone lysine demethylases, the Tet family of enzymes, and prolyl hydroxylase 2 (PHD2), which regulates stability of the transcription factor Hypoxia Inducible Factor α (HIF1α) (Williams et al., 2022, Chowdhury et al., 2011, Xu et al., 2011, Tyrakis et al., 2016, Intlekofer et al., 2017, Brinkley et al., 2020). Therefore, ectopic L-2HG accumulation can broadly disrupt mitochondrial metabolism, gene expression programs, chromatin architecture, and cell fate decisions (Intlekofer et al., 2017, Brinkley et al., 2020, Williams et al., 2022, Shelar et al., 2018, Taub et al., 2022, Ma et al., 2017, Tyrakis et al., 2016).

The ability of L-2HG to disrupt cellular processes forces animal cells to tightly control L-2HG metabolism. In this regard, the mitochondrial enzyme L-2-hydroxyglutarate dehydrogenase (L2hgdh) serves a key role in directly regulating L-2HG abundance by irreversibly converting L-2HG to αKG in an FAD-dependent manner (Rzem et al., 2004, Yang et al., 2023). The importance of this enzymatic reaction is evident by disease phenotypes caused by loss-of-function mutations in *L2hgdh* genes. Loss of L2hgdh activity in dogs, cats, and humans results in a rare disease known as L-2-hydroxyglutaric aciduria, which is characterized by developmental delays, seizures and other neurological symptoms, and in severe cases, childhood demise (Rzem et al., 2004, Rzem et al., 2007, Kranendijk et al., 2012, Christen et al., 2021, Christen et al., 2023, Sanchez-Masian et al., 2012, Farias et al., 2012). Similarly, *L2hgdh* mutant mice display both neurological pathologies and significant alterations in mitochondrial metabolism (Rzem et al., 2015, Ma et al., 2017, Brinkley et al., 2020). L-2HG also acts as an oncometabolite in the context of clear cell renal cell carcinomas (ccRCCs), where *L2hgdh* loss-of-function mutations induce ectopic L-2HG accumulation and lead to changes in DNA methylation, RNA methylation, mitochondrial metabolism, and amino acid synthesis. As a result, loss of L2hgdh activity within ccRCC cells promotes tumor growth and metastasis (Shim et al., 2014, Shelar et al., 2018, Brinkley et al., 2020, Taub et al., 2022).

While aberrant L-2HG accumulation is associated with several human diseases, mounting evidence indicates that L-2HG is also a product of normal cellular metabolism. For example, cultured cells accumulate L-2HG in response to hypoxia, low cellular pH, elevated NADH levels, and mitochondrial disruptions, suggesting that L-2HG production is part of the normal cellular response to oxidative stress (Intlekofer et al., 2015, Oldham et al., 2015, Mahmoudzadeh et al., 2020, Hunt et al., 2019, Li et al., 2020, Mullen et al., 2014, Nadtochiy et al., 2016, Reinecke et al., 2012). These links between L-2HG accumulation and cellular redox balance is further emphasized by the fact that L-2HG is synthesized from αKG by noncanonical activity of Lactate dehydrogenase (Ldh) and Malate dehydrogenase (Mdh) – two enzymes whose activity is intimately connected with cellular redox balance (Struys et al., 2007, Intlekofer et al., 2017, Intlekofer et al., 2015, Oldham et al., 2015, Rzem et al., 2007, Li et al., 2017, Teng et al., 2016). Finally, under certain circumstances, L-2HG appears capable of stabilizing HIF1α via inhibition of PHD2 (Intlekofer et al., 2017, Williams et al., 2022), thus highlighting a possible mechanism by which L-2HG production coordinates gene expression programs with changes in oxygen availability and cellular redox balance.

The importance of L-2HG metabolism is also becoming evident at the organismal level. A recent mouse study revealed that L2hgdh inhibition protects cardiac cells against ischemia-reperfusion injury (He et al., 2022), supporting the hypothesis that L2hgdh, and by extension L-2HG, plays an important and underappreciated role in the cellular response to oxidative stress. Similarly, L-2HG accumulates in CD8+ T-cells upon T-cell receptor activation and in macrophages stimulated with LPS (Tyrakis et al., 2016, Williams et al., 2022), with the accumulation of L-2HG in both cell types associated with HIF1α activation. These observations indicate that L-2HG metabolism serves an endogenous role in healthy tissues and suggest that additional studies of L-2HG and L2hgdh *in vivo* would lead to a better understanding of how ectopic L-2HG accumulation drives human disease.

*Drosophila* has emerged as an ideal system for studying L-2HG metabolism due to the facts that (i) the mechanisms that regulate L-2HG production and accumulation are conserved between flies and mammals (Yang et al., 2023, Li et al., 2017, Li et al., 2018), and (ii) *Drosophila* larvae accumulate millimolar levels of L-2HG during larval (juvenile) development in a controlled and highly predictable manner (Li et al., 2017).

Moreover, adult flies, like mammalian cells, generate high concentrations of L-2HG in response to hypoxia and oxidative stress (Mahmoudzadeh et al., 2020, Li et al., 2020, Hunt et al., 2019). These observations establish the fly as the only genetic model where L-2HG and L2hgdh functions can be studied *in vivo* with high levels of control and precision. Here we exploited this genetic system to demonstrate that renal L2hgdh activity is essential for maintaining mitochondrial metabolism and ensuring successful recovery from hypoxia.

Using previously described *L2hgdh* mutants (Li et al., 2017), we examined the effects of excess L-2HG accumulation on *Drosophila* growth, maturation, and metabolism. Our studies revealed that *L2hgdh* mutants exhibit few phenotypes under standard culture conditions but are hypersensitive to hypoxia. However, *L2hgdh* mutants do not die as the result of reduced oxygen availability, as nearly all mutant animals survive the hypoxia exposure; rather, *L2hgdh* mutants die during the reoxygenation phase with severe defects in mitochondrial metabolism. Subsequent tissue-specific studies revealed that the renal system (i.e., Malpighian tubules; MTs) is a key site of L2hgdh activity during hypoxia exposure, as expression of an *L2hgdh* transgene specifically within principal cells (PCs) of the MTs ensures successful recovery and rescues hypoxia-induced mitochondrial dysfunction. Finally, we demonstrated that L2hgdh serves an essential role in PCs even under normoxic conditions, as *L2hgdh* mutant PCs not only display reduced mitochondrial membrane potential and elevated ROS accumulation, but also exhibit aberrant growth upon Egfr activation. Overall, our studies reveal an ancient and conserved role for L2hgdh activity in the renal system and hints at a model in which this enzyme regulates L-2HG pool size as a means of coordinating mitochondrial flux with oxygen availability.

## RESULTS

### L2hgdh is required for recovery from hypoxia

To better understand the *in vivo* functions of L-2HG and L2hgdh (Figure 1A), we examined *Drosophila L2hgdh* mutants for defects in development, gene expression, and metabolism. Consistent with earlier studies (Li et al., 2017), we found that *L2hgdh^12/14^* mutant adult males accumulate very high L-2HG levels but exhibit no obvious differences in developmental timing, body mass, climbing behavior, or lifespan (Figure 1B, S1A-C, S2A,B). RNAseq analysis of *L2hgdh^12/14^*mutants also revealed modest changes in gene expression, 119 genes were up-regulated and 240 genes were down-regulated (Figure S3, Table S1, S2). Notably, we found no indication of enhanced HIF1α/Sima signaling or hypoxia-related signaling, as expression levels of *Ldh* (Wang et al., 2016), *Hph* (also known as *fatiga*; FBgn0264785)(Acevedo et al., 2010), and *Thor* (Barretto et al., 2020) were unchanged in *L2hgdh* mutants (Table S1, S2). *L2hgdh* mutations also had little effect on the expression of genes associated with metabolism, with only 44 metabolic genes exhibiting significantly altered mRNA transcript levels, none of which were associated with glycolysis, the TCA cycle, or the electron transport chain (Table S3). In fact, Pathway, Network and Gene-set Enrichment Analysis (PANGEA) analysis revealed only four GO categories that were significantly enriched among those genes that were either up- or down-regulated in *L2hgdh* mutants (Table S4)– sensory perception (GO:0007600), immune response (GO:0006955), cuticle development (GO:0042335), and response to external stimulus (GO:0009605).

**Figure 1.**
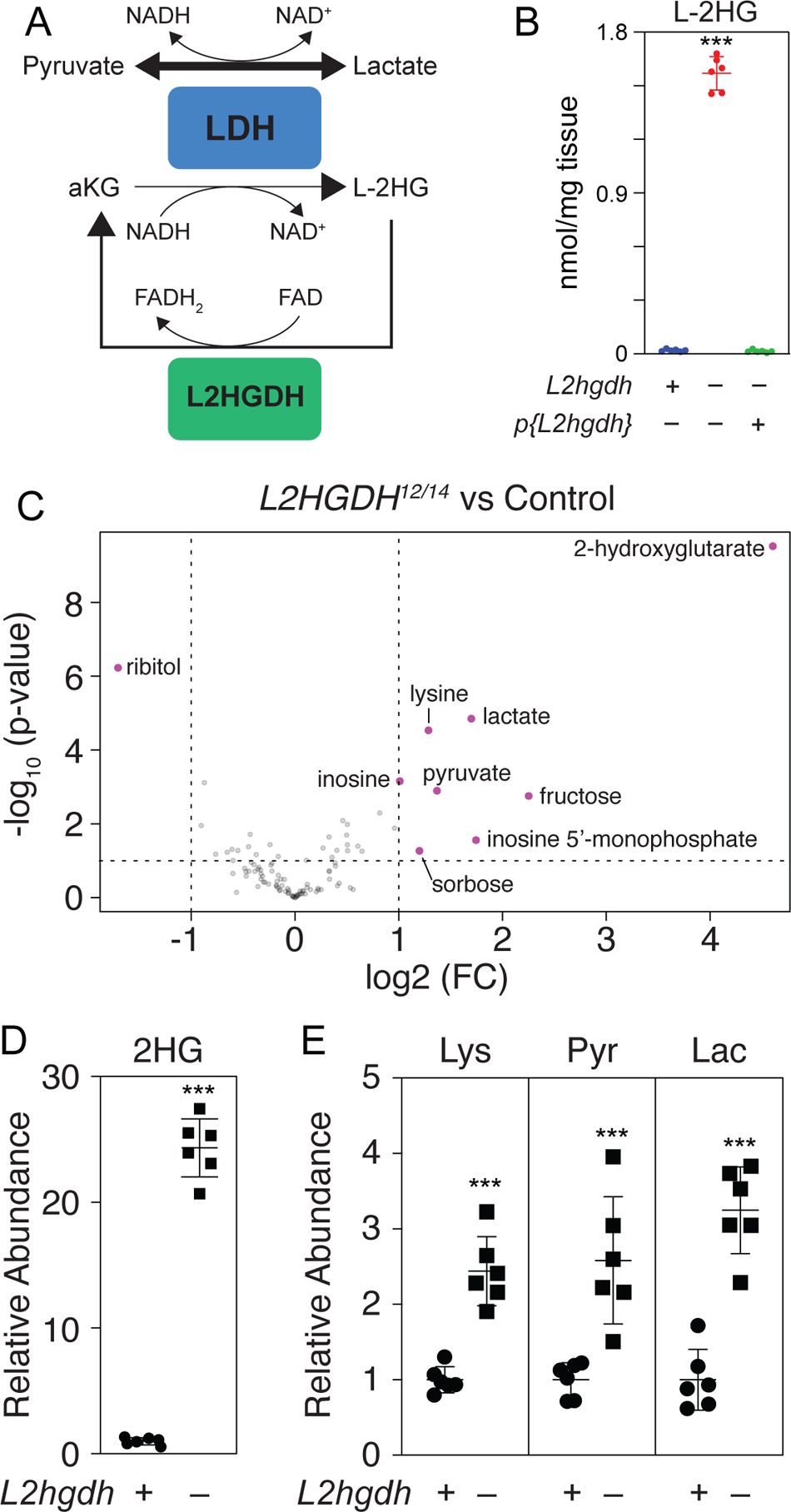
Metabolomic analysis of *L2hgdh* mutant adult males. (A) A schematic diagram representing the enzymatic reactions controlling L-2HG accumulation. Abbreviations are as follows: Lactate Dehydrogenase (LDH), L-2-hydroxyglutarate dehydrogenase (L2HGDH), L-2-hydroxyglutarate (L-2HG), α-ketoglutarate (αKG). (B) L-2HG levels were quantified in *L2hgdh^14/+^* controls, *L2hgdh^12/14^* mutants, and *L2hgdh^12/14^; p{L2hgdh}* rescued animals. (C-E) Data from GC-MS metabolomic analysis comparing *L2hgdh^14/+^* controls and *L2hgdh^12/14^* mutants analyzed with Metaboanalyst 5.0. (C) A volcano plot highlighting metabolites that exhibited a >1.5-fold change and *P* < 0.01. The relative abundance of (D) 2HG and (E) lysine (Lys), pyruvate (Pyr), and lactate (Lac) in *L2hgdh^14/+^* controls and *L2hgdh^12/14^* mutants presented as a scatter plot with the center line representing the mean and error bars representing the standard deviation. Genotype contrasts were performed with the Mann-Whitney test. *\*\*\*P*<0.001.

Considering that L-2HG and L2hgdh also regulate the immune response in mammals (Tyrakis et al., 2016, Williams et al., 2022), this finding is of considerable interest and will be the focus of future studies. Although ectopic L-2HG accumulation has little effect on *Drosophila* development, lifespan, or gene expression, a semi-targeted metabolomic analysis revealed that *L2hgdh^12/14^* mutant males displayed an unexpected metabolic profile when compared with *L2hgdh^14/+^* heterozygous controls (Figure 1C, S4, Table S5). While both 2HG and lysine were elevated in *L2hgdh^12/14^* mutants (Figure 1D,E), a result consistent with previous studies in mammals (Brinkley et al., 2020; Rzem *et al*., 2015), we also observed an unexpected increase in both lactate and pyruvate (Figure 1E). Subsequent GC-MS studies confirmed that *L2hgdh^12/14^* mutant males accumulate excess 2HG, lysine, and lactate in both whole animal extracts and hemolymph when compared with either the heterozygous control or a *L2hgdh^12/14^; p{L2hgdh}* rescue line (Figure S5A-D). Moreover, this increase in lactate accumulation was independent of changes in either *Ldh* gene expression or Ldh enzyme activity (Table S1, Figure S6). We also note that although pyruvate levels were inconsistently altered across these GC-MS-based validation experiments (Figure S5C), subsequent LC-MS-based studies described below revealed consistent increases in pyruvate levels. Overall, our results suggest that loss of L2hgdh activity shifts the balance of cellular metabolism towards a more glycolytic state in a manner that is independent of gene expression.

The presence of elevated lactate levels within *Drosophila L2hgdh* mutants suggests that loss of L2hgdh activity disrupts oxidative metabolism. We tested this possibility by asking if *L2hgdh* mutants are sensitive to hypoxia exposure (1% O_2_). Indeed, when compared with *L2hgdh^14/+^* heterozygous controls and a *L2hgdh^12/14^; p{L2hgdh}* rescue line, a significantly higher percentage of *L2hgdh^12/14^* mutants died following 6 hr, 12 hr, and 24 hr hypoxia treatment, with less than 25% of animals surviving the 12 hr and 24 hr exposures (Figure 2A-C). While quantifying survivorship, however, we noticed that greater than 90% of *L2hgdh^12/14^* mutant animals responded to physical touch immediately following removal from the hypoxia chamber (Figure 2D), indicating that *L2hgdh* mutants survive hypoxia but expire during the reoxygenation phase. Notably, this failure to recover from hypoxia exposure correlates with significantly elevated L-2HG levels in *L2hgdh* mutants, both immediately following 12 hr hypoxia exposure as well as 1 hr post-recovery (Figure 2E). Together, our findings indicate that L2hgdh limits L-2HG accumulation in hypoxia-exposed animals and ensures successful recovery.

**Figure 2.**
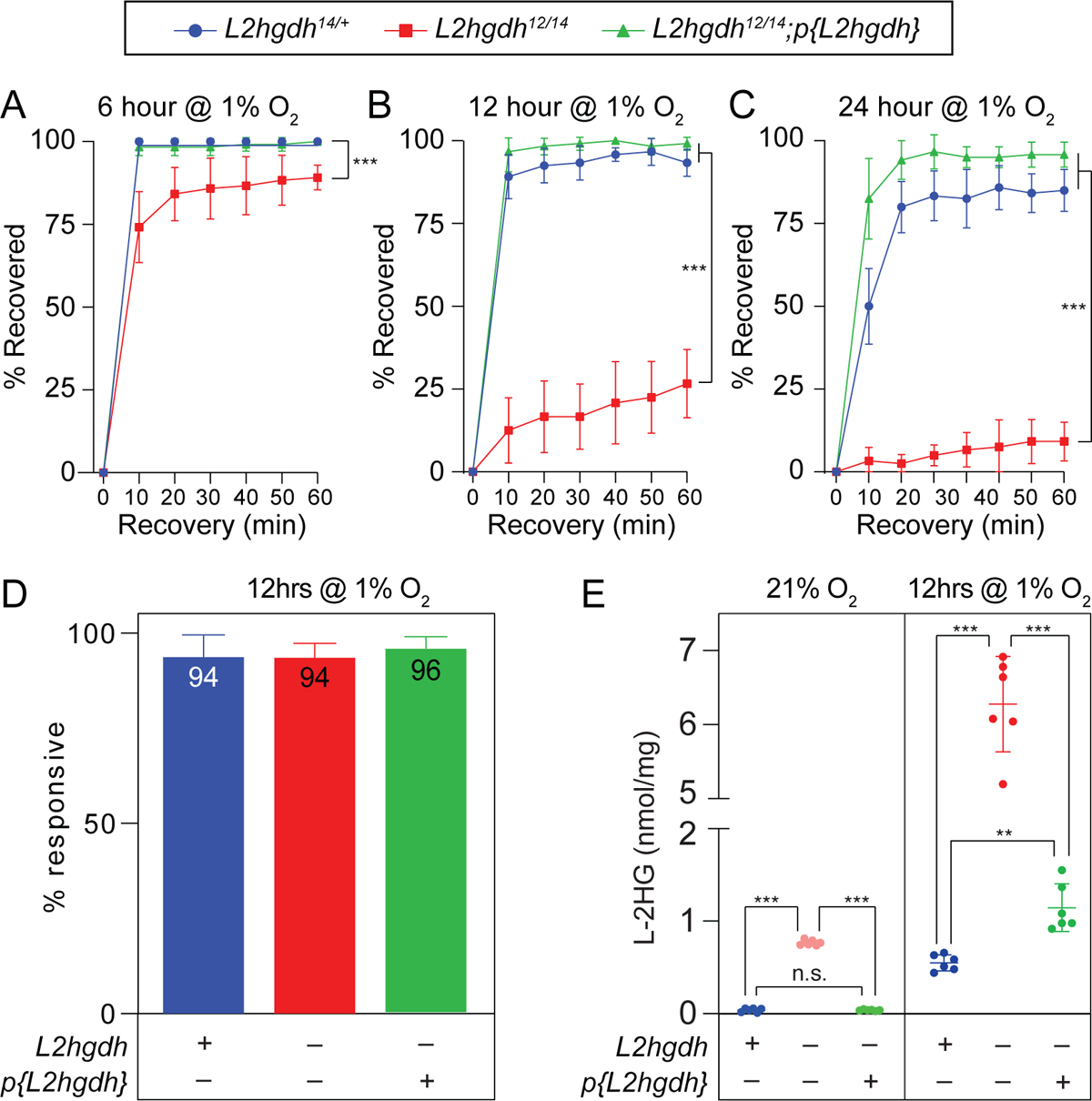
*L2hgdh* adult mutant males are hypersensitive to hypoxia exposure. *L2hgdh^14/+^* controls, *L2hgdh^12/14^* mutants, and *L2hgdh^12/14^; p{L2hgdh}* rescued adult male flies were exposed to 1% O_2_ for (A) 6 hrs, (B) 12 hrs, or (C) 24 hrs and monitored for recovery over a 90-minute period. n=6 vials containing 10 adult male flies for each genotype. For (A-C), the data were subject to repeated measures ANOVA with a Geisser-Greenhouse correction, followed by a Holm-Sidak’s multiple comparison test \*\*\**P*>0.001. (D) Flies were removed from the hypoxia chamber and the ability of individual animals to respond to mechanical stimulation was assessed. (E) L-2HG levels in normoxic and hypoxic conditions across genotypes. For (D,E), data analyzed using Brown-Forsythe ANOVA test followed by a Dunnett’s multiple comparison test. *\*\*P>*0.01. \*\*\**P*>0.001.

### Mitochondrial metabolism is disrupted in hypoxia-treated *L2hgdh* mutants

To further explore the function of L2hgdh in the hypoxia response, we used semi-untargeted LC-MS-based metabolomics to examine *L2hgdh* mutants and the two control strains at three timepoints: (i) immediately following a 12 hr incubation in normoxia (Norm), (ii) immediately following a 12 hr hypoxia exposure (H+0 hr), and (iii) 12 hrs of hypoxia followed by 1 hr of normoxia (H+1 hr; note that we only assayed living flies; Table S6). Principal component analysis (PCA) revealed that the normoxic metabolome of *L2hgdh* mutants was similar to that of the *L2hgdh^14/+^* heterozygous controls and the *L2hgdh^12/14^; p{L2hgdh}* rescue line (Figure 3A). However, hypoxia (PC1) induced significant metabolic changes in *L2hgdh* mutant males relative to the controls, and these differences persisted at the H+1 hr timepoint (Figure 3A). Moreover, while the H+0 hr and H+1 hr metabolomes of both control groups separate with no overlap of their confidence intervals (Figure 3A), the H and H+1 *L2hgdh* mutants sample groups segregate from the controls along PC2 and completely overlap one another, suggesting that L2hgdh is essential for the metabolome to recover from hypoxia treatment. Consistent with the PCA analysis, hierarchical clustering reveals that the H+0 hr and H+1 hr *L2hgdh* mutant samples share a lowest common ancestor (Figure S7).

**Figure 3.**
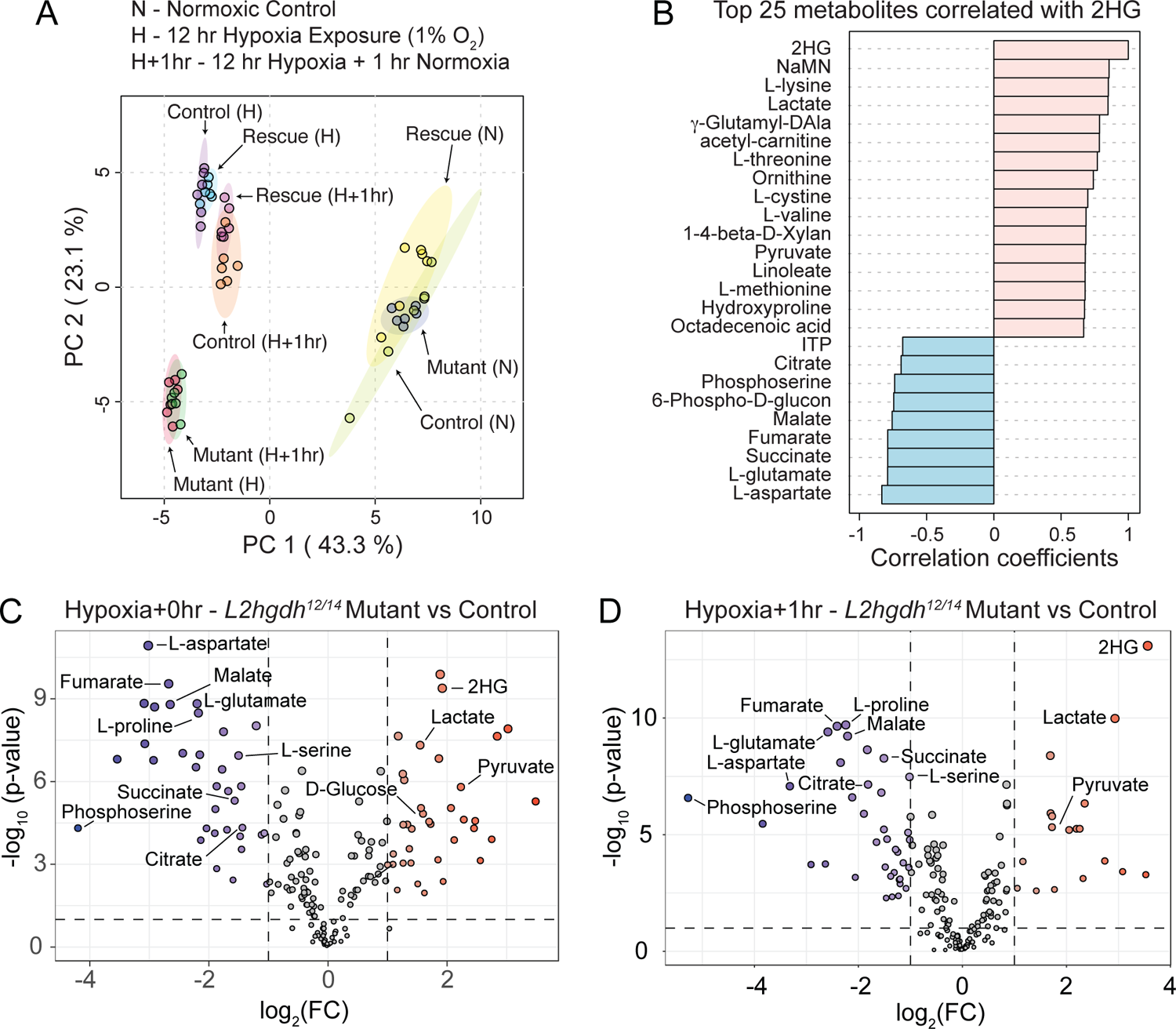
Metabolomic analysis of hypoxia-exposed *L2hgdh* mutants. *L2hgdh^14/+^*controls, *L2hgdh^12/14^* mutants, and *L2hgdh^12/14^; p{L2hgdh}* rescued adult male flies were analyzed using a semi-targeted LC-based metabolomics following either (i) a 12 hr incubation in 1% O_2_ (H), (ii) a 12 hr incubation in 1% O_2_ followed by a 1 hr recovery in normoxic conditions (H+1 hr), (iii) a 12 hr incubation in normoxic conditions. (A) PCA analysis of the nine groups of analyzed samples. Each genotype abbreviated as follows: *L2hgdh^14/+^* (Control), *L2hgdh^12/14^* (Mutant), *L2hgdh^12/14^; p{L2hgdh}* (Rescue). (B) Top 25 metabolites correlated with 2HG levels across all samples. (C) A volcano plot highlighting metabolites that exhibited a >2-fold change and *P*<0.01 in control vs. mutant samples immediately following the 12 hr hypoxia exposure. (D) As in (C) but following a 12 hr hypoxia exposure and 1 hr recovery. For all genotypes and conditions, n=6 biological replicates containing 20 adult males. All panels generated with Metaboanalyst 5.0.

The control and rescue strains, however, primarily segregate by recovery time, with the H+0 hr and H+1 hr samples from these genotypes defining distinct subclusters (Figure S7). Overall, our analyses demonstrate that the *L2hgdh* mutant metabolome is significantly disrupted by hypoxia exposure and indicate that L2hgdh activity is essential during the reoxygenation phase for to reestablish normal oxidative metabolism.

To determine what factors drive the metabolomic differences between the control, rescue, and mutant strains, we used a correlation analysis to identify metabolites that are altered in response to elevated 2HG levels (note that L-2HG represents the bulk of the adult 2HG pool upon hypoxia treatment, see Figure 2E). This approach revealed that L-2HG levels positively correlate with the abundance of pyruvate and lactate, among others (Figure 3B). In contrast, concentrations of the TCA cycle metabolites citrate, succinate, fumarate, and malate, as well as the anaplerotic amino acids glutamate, proline, and aspartate, displayed inversely proportional relationships with L-2HG abundance (Figure 3B).

The importance of these metabolomic results become apparent upon closer examination of the datasets. As expected, hypoxia induced 2HG and lactate accumulation in all three strains; however, hypoxia-exposed *L2hgdh* mutants display a significantly larger increase in the abundance of both metabolites (Figure 3C,D, and S8). Moreover, although the control and rescue strain contained similar amounts of glucose and pyruvate across all three timepoints, hypoxia-treated *L2hgdh* mutants harbored elevated amounts of both molecules at H+0 and pyruvate levels remained elevated at H+1 (Figure 3D, S8). Similarly, while levels of citrate, succinate, fumarate, and malate, as well as glutamate, proline, and aspartate, remained at near constant levels in control strains regardless of timepoint, *L2hgdh* mutants displayed significant decreases in all of these molecules at both the H+0 hr and H+1 hr timepoints (Figure 3C,D, S9A-D, S10A-C). Intriguingly, we also noted that while the control strains accumulate higher levels of serine following hypoxia treatment, *L2hgdh* mutants do not (Figure S10D). This result is notable considering that raised L-2HG (due to loss of L2HGDH expression) also disrupt serine metabolism on renal cancer cells (Kundu et al., 2024).

We confirmed our metabolomic results using GC-MS to measure the relative abundance of specific metabolites in glycolysis, the TCA cycle, as well as a subset of amino acids in the control, mutant, and rescue strain. For this analysis, we assayed the three timepoints described above as well as a fourth timepoint, 12 hours of hypoxia followed by 12 hours of normoxia (H+12). The resulting data mirrored those generated in our LC-MS study – in all three strains, hypoxia induced elevated levels of 2HG and lactate, with *L2hgdh* mutant adults displaying significantly increased quantities of 2HG and lactate at both the H+0 and H+1 timepoints (Figure 4A,B,D,E). We also observed elevated 2HG levels in *L2hgdh* mutant at the H+12 timepoint (Figure 4C), but the relative abundance of L-2HG at this timepoint was less than that observed at H+0 and H+1 (Figure 4A-C). Similarly, lactate levels returned to normal in H+12 *L2hgdh* mutants (Figure 4F).

**Figure 4.**
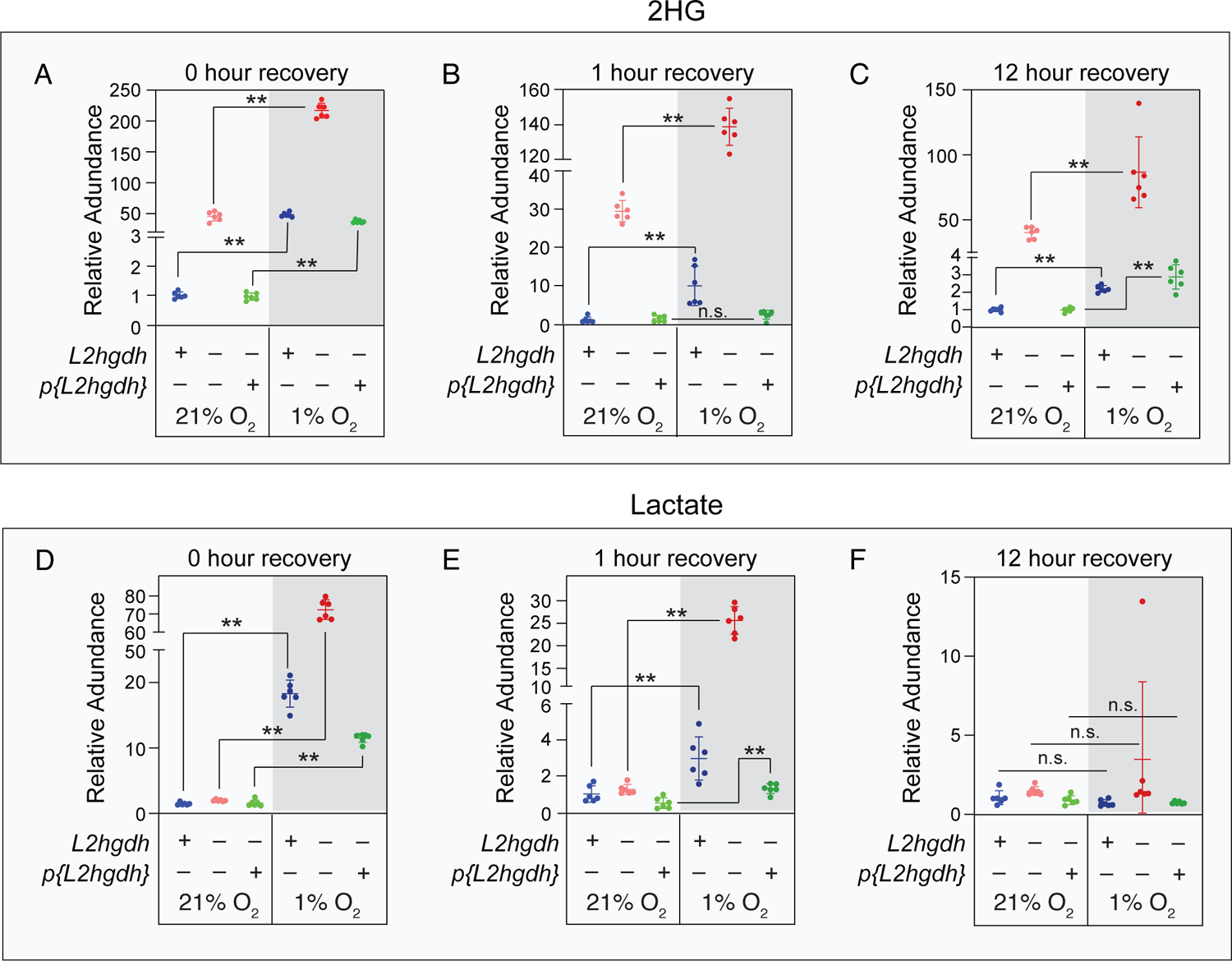
2HG and lactate levels remain elevated in *L2hgdh* mutants following hypoxia exposure. The relative abundance of (A-C) 2-hydroxyglutarate (2HG) and (D-F) lactate were measured using GC-MS in *L2hgdh^14/+^* controls, *L2hgdh^12/14^* mutants, and *L2hgdh^12/14^; p{L2hgdh}* rescued adult male flies following either (i) a 12 hr incubation in 1% O_2_ (H), (ii) a 12 hr incubation in 1% O_2_ followed by a 1 hr recovery in normoxic conditions (H+1 hr), (iii) a 12 hr incubation in 1% O_2_ followed by a 12 hr recovery in normoxic conditions (H+12 hr). A set of normoxic control samples were collected at each timepoint. For all genotypes and conditions, n=6 biological replicates containing 20 adult males. Data in all panels was statistically analyzed by comparing the normoxic and hypoxic samples for each genotype using a Mann-Whitney test. *\*\*P>*0.01. n.s. (not significant).

Analysis of TCA cycle intermediates similarly validated our initial metabolomic study. While αKG, succinate, fumarate, and malate levels were decreased in the hypoxia-treated control strains at H+0, the *L2hgdh* mutants displayed a notably larger decrease in nearly all TCA cycle intermediates at this timepoint (Figure 5A). Moreover, while levels of TCA cycle intermediates in the H+1 control strain had recovered to those observed in untreated samples, H+1 mutant samples contained significantly lower levels of citrate, succinate, fumarate, and malate (Figure 5B). At the H+12 timepoints, however, nearly all TCA cycle metabolites either returned to levels seen in the normoxic controls or exhibited an unexpected increase – a trend observed in all three genotypes (Figure 5C). Overall, our results suggest that L2hgdh limits the amount of L-2HG that accumulates during hypoxia exposure, and in its the absence, *L2hgdh* mutants fail to strike a proper balance between glycolytic and mitochondrial metabolism, resulting in death during reoxygenation.

**Figure 5.**
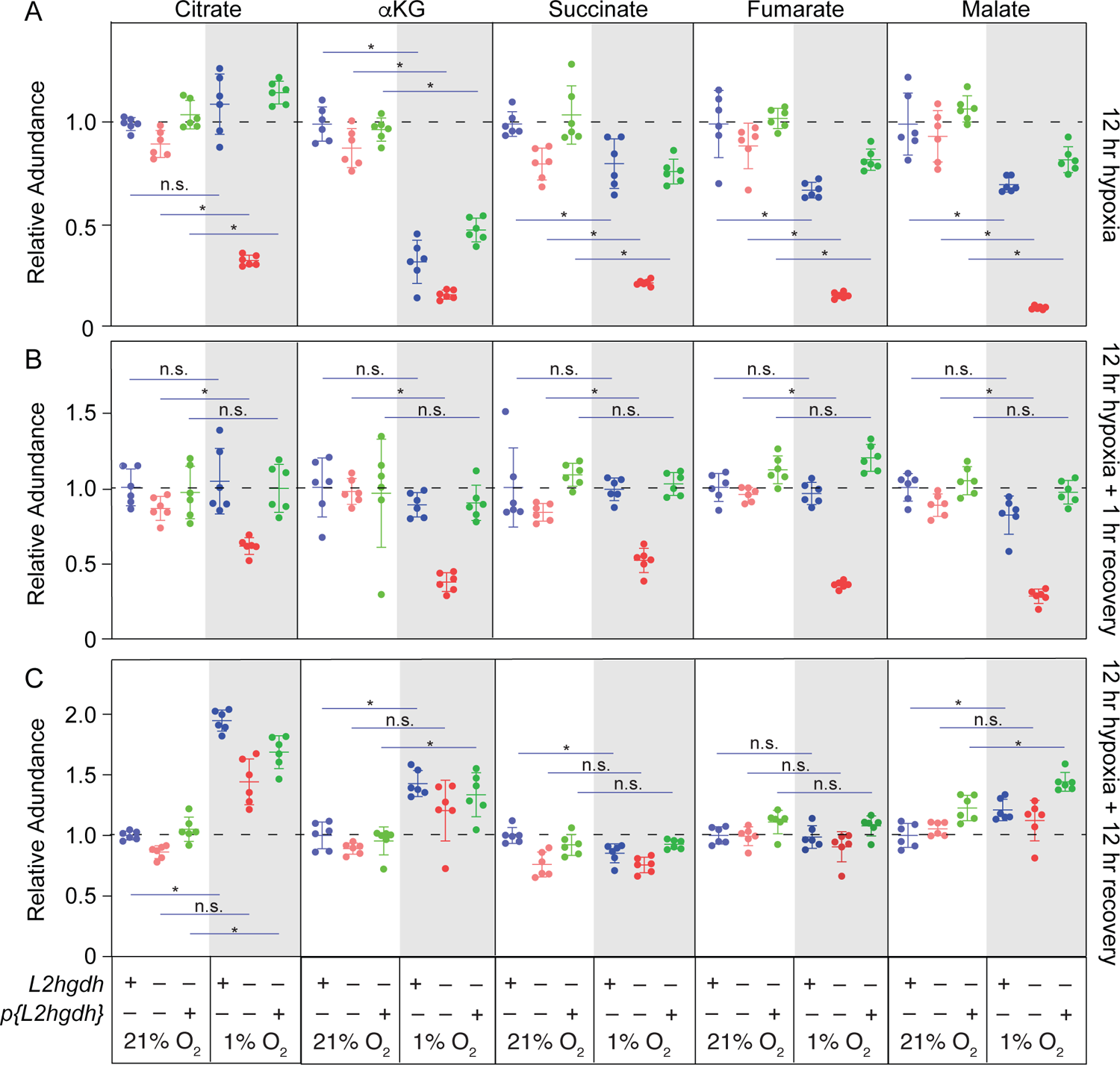
*L2hgdh* mutants display defects in mitochondrial metabolism during reoxygenation. (A-C) The relative abundance of the TCA cycle metabolites citrate, α-ketoglutarate (αKG), succinate, fumarate, and malate was measured using GC-MS in *L2hgdh^12/14^* mutants, and *L2hgdh^12/14^; p{L2hgdh}* rescued adult male flies following either (i) a 12 hr incubation in 1% O_2_ (H), (ii) a 12 hr incubation in 1% O_2_ followed by a 1 hr recovery in normoxic conditions (H+1 hr), (iii) a 12 hr incubation in 1% O_2_ followed by a 12 hr recovery in normoxic conditions (H+12 hr). A set of normoxic control samples were collected at each timepoint. Data in all panels were analyzed by comparing the normoxic and hypoxic samples for each genotype using the Mann-Whitney test. *\*\*P>*0.01. n.s. (not significant).

### Hif1α target genes are properly regulated in hypoxia-treated *L2hgdh* mutants

As a complement to the metabolomic studies, we compared gene expression in *L2hgdh* mutants with *L2hgdh^14/+^* heterozygous control animals at the same four timepoints used in our targeted metabolomic assays (as in Figures 4, 5). Both strains responded to changes in oxygen availability by activating similar gene expression programs, with mutant and control samples exhibiting similar numbers of up- and down-regulated genes (Table 1, Tables S7-S12). PCA analysis of differentially expressed genes across the genotypes and timecourse placed the H+0 and H+1 datasets at similar positions on the PC1 axis (Figure 6A), which is strongly correlated with treatment condition (hypoxia) and a combination of treatment and genotype (Figure S11A). In addition, the mutant and control samples at each timepoint segregate along the PC2 axis, which is largely attributable to genotype (Figure S11A). Interestingly, we note that the H+12 samples locate midway between the normoxic and H+0/H+1 samples along the PC1 axis (Figure 6A), suggesting that gene expression in both control and mutant animals was returning to normoxic levels by 12h. These patterns are also reflected in a hierarchical clustering analysis, which revealed that the normoxic and H+12 samples grouped together under a lowest common parent node, as did the H+0 and H+1 samples (Figure 6B). Overall, these observations suggest that control and *L2hgdh* mutants mount similar transcriptional responses to hypoxia and reoxygenation.

**Figure 6.**
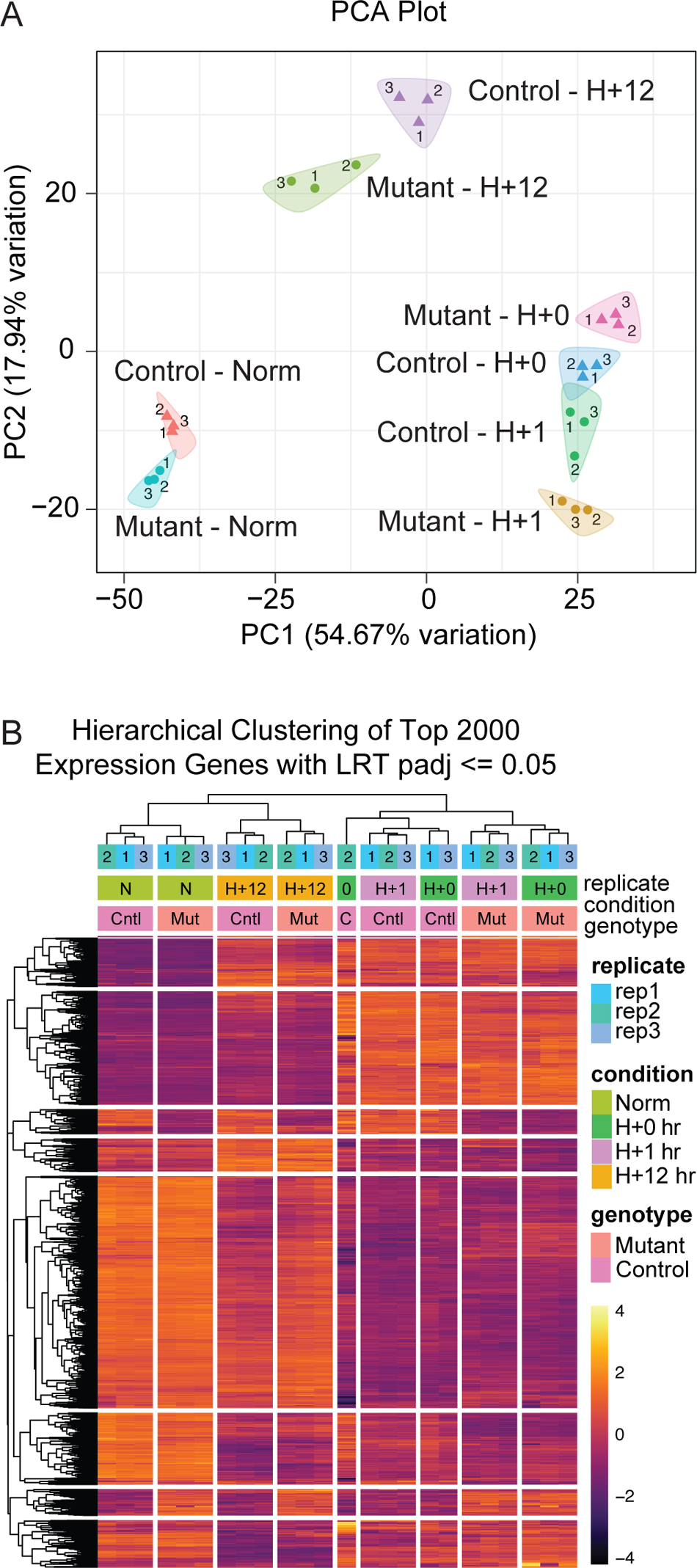
RNAseq analysis of hypoxia exposed *L2hgdh* mutants. (A) PCA biplot of mutant and control normoxic, hypoxic, and recovered replicates, PC1 vs. PC2. (B) Complete hierarchical clustering of the top 2,000 genes by variance with LRT adjusted p-value < 0.05 across all samples. Upper bars denote sample annotations.

**Table 1.** Number of genes significantly changed in hypoxia-treated *L2hgdh^14/+^* controls and *L2hgdh^12/14^* mutants samples relative to normoxia control samples.

| Condition | Normoxia |  | Hypoxia + 0 hr recovery |  |  |  | Hypoxia + 1 hr recovery |  |  |  | Hypoxia + 12 hr recovery |  |  |  |
| --- | --- | --- | --- | --- | --- | --- | --- | --- | --- | --- | --- | --- | --- | --- |
| Genotype | Mutant |  | Control |  | Mutant |  | Control |  | Mutant |  | Control |  | Mutant |  |
| Direction of gene expression | Up | Down | Up | Down | Up | Down | Up | Down | Up | Down | Up | Down | Up | Down |
| # genes significantly changed | 47 | 57 | 636 | 1139 | 633 | 1070 | 626 | 1288 | 586 | 1099 | 310 | 747 | 258 | 349 |

A closer examination of the RNA-seq data revealed that not only were fewer than 260 genes significantly altered in the mutant strain at any of the four timepoints (Table S8), but the Hif1α/sima target genes *Ldh* and *Hph* were also expressed at similar levels in mutant and control animals throughout the timecourse (Figure S11B-D). PANGEA analysis of the RNAseq data revealed that, under both normoxia and hypoxia, the most significantly altered Gene Set present at all timepoints was “Toll and Imd signaling pathway” (KEGG path:map04624; Table S13), again highlighting the link between L-2HG metabolism and the innate immune response. There was, however, no enrichment for gene categories or pathways related to glycolysis, mitochondrial metabolism, or HIF1α/sima signaling (Table S13). These results indicate that the hypoxia induced metabolic defects observed in *L2hgdh* mutants stem from disruption of mitochondrial metabolism, not a defect in HIF1α/sima signaling.

### Renal L2hgdh activity ensures recovery from hypoxia treatment

To better understand why *L2hgdh* mutants exhibit defects in mitochondrial metabolism and hypoxia sensitivity, we shifted from whole-animal analyses to tissue-specific studies of the renal and nervous systems – tissues that both express significant levels of L2hgdh transcripts and exhibit the most dramatic phenotypes in *L2hgdh* mutant mammals (Li et al., 2022, Shim et al., 2014, Rzem et al., 2015, Rzem et al., 2004). For our initial tissue-specific studies, we exposed *L2hgdh^12/14^* mutants that expressed a *UAS-L2hgdh* transgene in either neurons (*elav-GAL4*)(Yannoni and White, 1997) or principal cells (PCs) of the Malpighian tubules (*C42-GAL4*)(Rosay et al., 1997) to 1% O_2_ for 12 hours. To our surprise, all *L2hgdh^12/14^; C42-GAL4 UAS-L2hgdh* animals (from here on referred to as *L2hgdh; C42-L2hgdh*) survived 12 hr hypoxia exposures (Figure 7A), while the negative control strains (i.e., *L2hgdh^12/14^; C42-GAL4/+* and *L2hgdh^12/14^; +/UAS-L2hgdh*) died at the same rate as *L2hgdh^12/14^* mutants (Figure 7A). In contrast, *L2hgdh^12/14^; elav-GAL4 +/+ UAS-L2hgdh* (now referred to as *L2hgdh; elav-L2hgdh*) died at the same rate as the negative control strains (Figure S12A).

**Figure 7.**
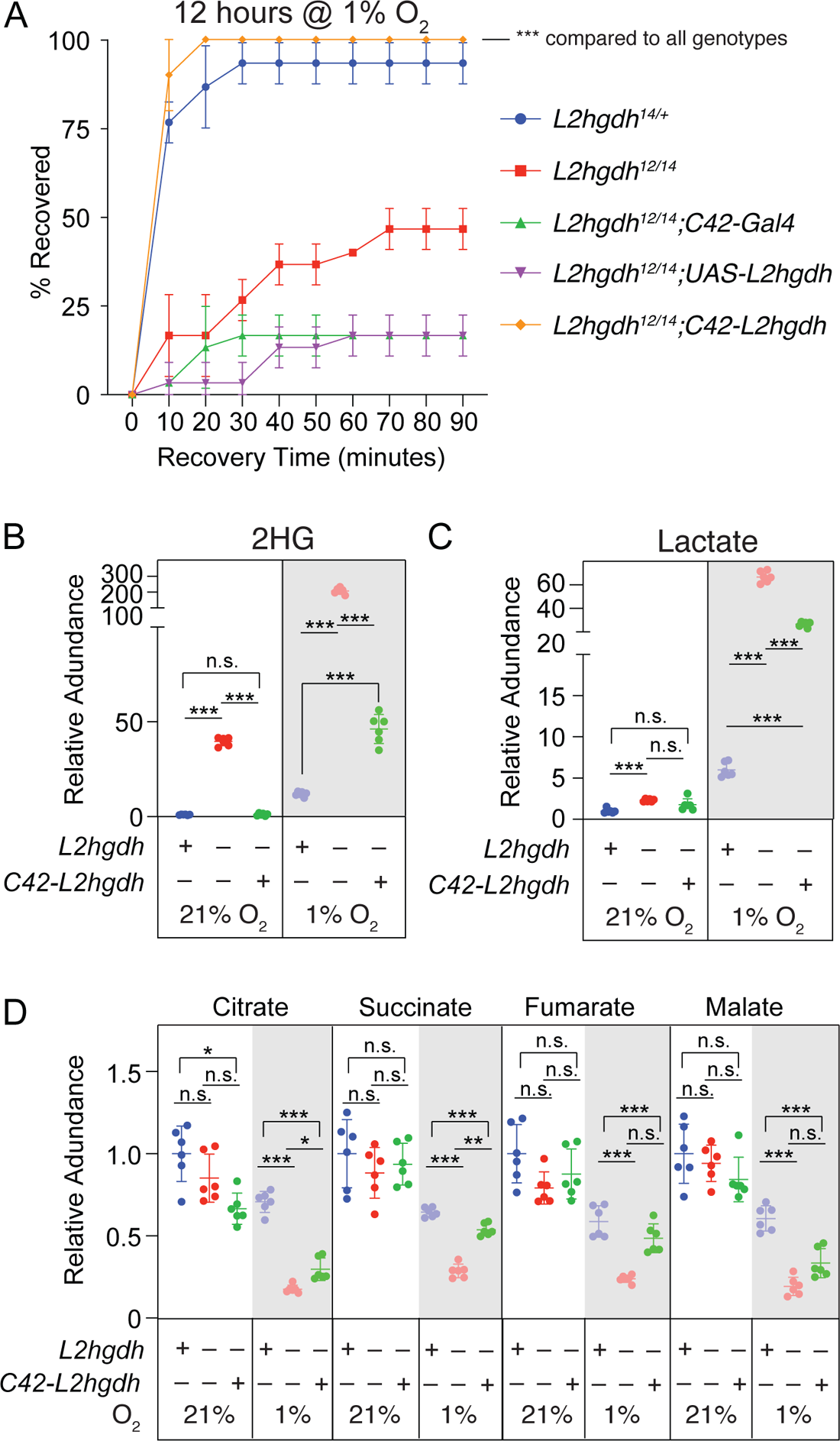
*L2hgdh* expression in the renal system is sufficient to support viability during reoxygenation. (A) Adult male flies were exposed to 1% O_2_ for 12 hours and monitored for recovery over the course of 90 minutes. n=6 vials containing 10 adult male flies for each genotype. Data analyzed using a repeated measures ANOVA with a Geisser-Greenhouse correction followed by a Holm-Sidak’s multiple comparison test. \*\*\**P*>0.001. n=3 vials with 10 adult male flies per vial. (B-D) GC-MS was used to measure (B) 2HG, (C) lactate, and (D) the TCA cycle intermediates citrate, succinate, fumarate, and malate in *L2hgdh^14/+^* controls, *L2hgdh^12/14^* mutants, and *L2hgdh^12/14^; C42-Gal4 UAS-L2hgdh* rescued animals. For (B-D), data analyzed using Brown-Forsythe ANOVA test followed by a Dunnett’s multiple comparison test. *\*\*P>*0.01. \*\*\**P*>0.001.

Beyond the viability phenotype, *C42-L2hgdh* expression in hypoxia-exposed *L2hgdh* mutants also partially restored whole body 2HG, lactate, citrate, and succinate levels (Figure 7B-D). Moreover, although fumarate and malate levels were not significantly rescued in *L2hgdh; C42-L2hgdh* animals when compared with mutants, the abundance of both metabolites trended higher in the rescued animals. In contrast, *L2hgdh; elav-L2hgdh* expression failed to significantly rescue the abundance of any of the assayed metabolites (Figure S12B-D). These findings indicate that L2hgdh activity within the renal system is sufficient to maintain systemic mitochondrial activity and enable successful recovery from hypoxia exposure.

### Renal L2hgdh activity controls L-2HG excretion and catabolism

The ability of *C42-L2hgdh* expression to rescue the *L2hgdh* mutant phenotypes raises the question of how renal L2hgdh activity can have such profound effects on organismal physiology and metabolism. A clue towards answering this question comes from the observation that urinary L-2HG levels are elevated in both *L2hgdh* KO mice and individuals suffering from L-2HG aciduria (Rzem et al., 2015, Ma et al., 2017, Duran et al., 1980, Kranendijk et al., 2012) (Figure S13). These observations highlight how the renal system responds to loss of L2hgdh activity by attempting to excrete excess L-2HG. Considering that the PCs are a key site of L2hgdh function, and that these cells are responsible for transport of organic metabolites (Cohen et al., 2020), we used two approaches to determine if the fly renal system also excretes L-2HG:

i. We used GC-MS to quantify L-2HG within the excreta of *L2hgdh* mutants. Consistent with previous mammalian observations, the excrement of *L2hgdh^12/14^* mutant male flies contained significantly elevated L-2HG levels when compared with the control strain (Figure 8B), indicating that flies, like mammals, excrete accumulated L-2HG upon loss of L2hgdh activity.
ii. We measured the secretion rate of isolated Malpighian tubules using a Ramsay assay to determine if loss of L2hgdh alters renal activity. Our analysis revealed that the secretion rate of *L2hgdh^12/14^*mutant MTs was significantly higher than that measured in either the control or rescue line (Figure 8A), suggesting that the elevated levels of L-2HG in *L2hgdh* mutants induce increased excretion.

**Figure 8.**
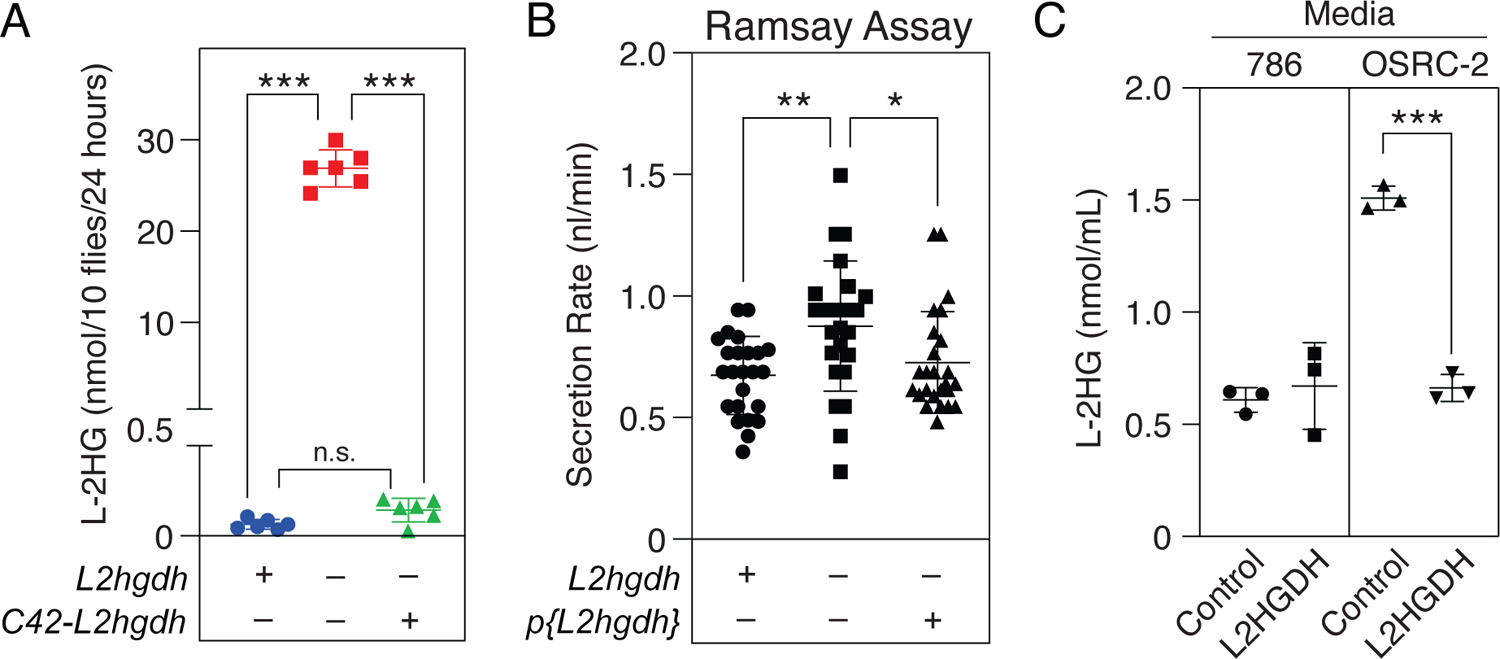
*L2hgdh* mutants exhibit higher rates of renal activity. (A) L-2HG levels quantified in the excreta of *L2hgdh^14/+^* controls, *L2hgdh^12/14^* mutants, and *L2hgdh^12/14^; C42-Gal4 UAS-L2hgdh* rescued animals. (B) A Ramsay assay was used to measure the secretion rate of MTs isolated from *L2hgdh^14/+^* controls, *L2hgdh^12/14^* mutants, and *L2hgdh^12/14^; p{L2hgdh}* rescued animals. (C) L-2HG levels were significantly elevated in the culture media of OSRC-2 cells when compared with 786 cells. L2hgdh transfection into OSRC-2 cells, but not 786 cells, significantly reduced L-2HG levels in media. Statistical analysis of all panels conducted using a Mann-Whitney test. *\*\*\*P*<0.001.

Our findings raise the question as to whether the primary purpose of renal L2hgdh activity is to promote L-2HG catabolism or excretion. If L2hgdh promotes L-2HG excretion, we would expect to see increased L-2HG levels in the excreta of *L2hgdh; C42-L2hgdh* compared with the mutant strain. In contrast, if L2hgdh modulates systemic L-2HG levels by converting L-2HG to αKG, then the excreta of *L2hgdh; C42-L2hgdh* should contain L-2HG levels comparable to the control strains. We observed the latter, as GC-MS analysis of excreta from the *L2hgdh; C42-L2hgdh* strain contained the same amount of L-2HG as the heterozygous controls (Figure 8B). This observation indicates that renal L2hgdh activity catabolizes the majority of the normoxic L-2HG pool while loss of this enzyme results in increased L-2HG excretion.

The ability of renal L2hgdh activity to influence both steady state L-2HG levels and MT secretion rate motivated us to determine if human L2hgdh also influences L-2HG excretion from the human renal carcinoma cell lines 786-O and OSRC-2. While previous studies demonstrated that L2hgdh activity regulates intracellular L-2HG accumulation within these cells, the influence of L2hgdh on L-2HG excretions is unknown. We found that while L-2HG concentration in the media of 786-O cells was similar between those transfected with either the control vector or the L2HGDH expressing plasmid, the media of OSRC-2 cells expressing the empty vector control contained significantly higher levels of L-2HG than the media of cells expressing L2HGDH (Figure 8C). Together, our findings suggest that L-2HG transport could play an important role in ccRCC.

### L2hgdh functions within PCs to maintain mitochondrial metabolism

Our results indicate that ectopic L-2HG accumulation inhibits mitochondrial metabolism and highlight the PCs as a key site of L2hgdh activity. To further explore the role of L2hgdh within these cells, we used the dye JC1, which stains both mitochondria (JC1 Green) and mitochondrial membrane potential (JC1 Red), to determine how loss of L2hgdh activity influences cell autonomous mitochondrial metabolism. Our study revealed that, even under normoxic conditions, staining of JC1 Red relative to JC1 Green was significantly reduced in the PCs of *L2hgdh* mutant MTs (Figure 9A and A’) when compared with *L2hgdh^14/+^*controls (Figure 9B and B’). Moreover, the decreased JC1 Red staining observed in *L2hgdh* mutant MTs was completely rescued by expression of a *UAS-L2hgdh* transgene using the PC-specific *C42-Gal4* driver (*C42-L2hgdh*; Figure 9C and C’). Unfortunately, we were unable to conduct these assays after hypoxia treatment due to the extreme fragility of hypoxia-exposed *L2hgdh* mutant MTs. Together, our observations suggest that even under normoxic conditions, L2hgdh functions within the PCs to promote mitochondrial metabolism in a cell autonomous manner.

**Figure 9.**
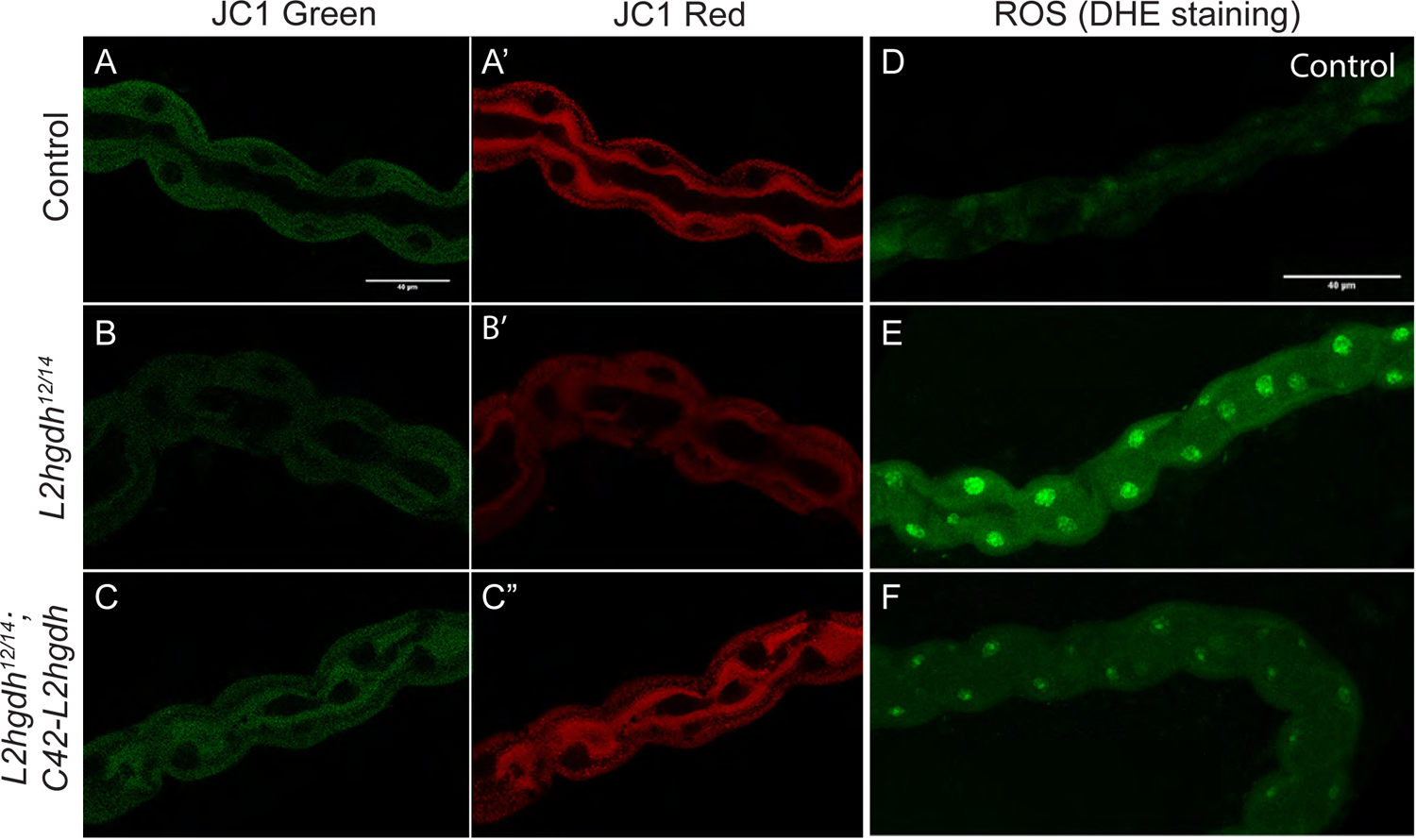
L2hgdh supports mitochondrial metabolism in PCs. MTs from *L2hgdh^14/+^* controls, *L2hgdh^12/14^* mutants, and *L2hgdh^12/14^; C42-Gal4 UAS-L2hgdh* rescued animals were stained using either (A-C) JC1 or (D-F) DHE. (A-C) While JC1 green was present at similar levels in PCs of all genotypes, (A’-C’) JC1 red staining was lower in *L2hgdh^12/14^* mutant principal cells as compared with either the control or *L2hgdh^12/14^; C42-Gal4 UAS-L2hgdh* rescue strain. (D,E) DHE staining of adult male MTs revealed that ROS levels were elevated in *L2hgdh^12/14^* mutant principal cells as compared with *L2hgdh^14/+^* controls. Meanwhile, (F) PCs of the *L2hgdh^12/14^; C42-Gal4 UAS-L2hgdh* rescue strain exhibit ROS levels comparable to those observed in (D) the control strain. 40 μm scale bar in (A) also applies to (B) and (C). 40 μm scale bar in (D) also applies to (E) and (F).

As a complement to these dye-based assays, we used RNA-seq to determine if the observed mitochondrial defects stem from changes in MT gene expression. Similar to the whole animal analyses, only 124 genes were up-regulated and 90 genes were down-regulated in *L2hgdh* mutant MTs when compared with the controls (Figure S14A, Table S14 and S15). We again observed no significant link between loss of L2hgdh activity and expression of genes associated with glycolysis, mitochondrial metabolism, or HIF1α signaling– one gene involved in glycolysis was upregulated in mutant MTs, and there were no significant changes in expression among those genes involved in the TCA cycle or electron transport chain (Figure S14B-D, Table S15). Further analysis using PANGEA revealed that the only metabolic pathways significantly altered in *L2hgdh* mutants were sulfur metabolism (<u>GO:0006790</u>) and cofactor metabolic processes (<u>GO:0051186</u>; Table S16). These enrichments, however, were driven entirely by changes in genes encoding glutathione S-transferases (GSTs; Table S16). Considering that *L2hgdh* mutant MTs exhibit elevated secretion rates while also experiencing compromised mitochondrial metabolism, increased GST expression could be indicative of elevated ROS production. Indeed, the MTs of *L2hgdh* mutants exhibited higher levels of ROS, as determined by DHE staining, when compared with controls (Figure 9D,E), and these metabolic phenotypes were rescued by expressing *C42-L2hgdh* in *L2hgdh* mutant PCs (Figure 9E). Thus, loss of L2hgdh expression within the PCs results in an aberrant metabolic state that leads to elevated ROS production.

### *L2hgdh* mutant MTs are sensitized towards aberrant tissue growth

The decreased mitochondrial activity and elevated ROS levels present within *L2hgdh* mutant MTs mirror the metabolic state of ccRCC tumors, which are highly glycolytic, display decreased mitochondrial respiration, and exhibit elevated ROS production (Courtney et al., 2018). Based on the metabolic phenotypes shared between the *Drosophila L2hgdh* mutant MTs and ccRCCs, as well as the fact that L2hgdh functions as a tumor suppressor in renal cells, we hypothesized that *L2hgdh* mutant MTs are sensitized toward aberrant growth upon manipulation of genes involved in ccRCC. We initially evaluated this possibility by expressing a *UAS-Vhl-RNAi* construct in the PCs of adult *L2hgdh* mutants using *C42-GAL4* under the control of *Gal80ts* (*L2hgdh; C42-Vhl-RNAi; Gal80ts*). Considering that *Vhl* is mutated in nearly 90% of ccRCCs (Gossage et al., 2015), we predicted that *Vhl-RNAi* would interact with *L2hgdh* mutations to induce synthetic MT phenotypes. The resulting genetic combination, however, was extremely sick and we were unable generate *L2hgdh; C42-Vhl-RNAi; Gal80ts* adult males, even when reared at the permissive temperature for Gal80ts.

While the few resulting adults showed no MT abnormalities (data not shown), these analyses inevitably reflect a survivor bias, as we hypothesize that only those animals with inefficient *Vhl* knockdown completed development. This observation demonstrates a genetic interaction between *L2hgdh* and *Vhl* that will be explored in future studies.

As an alternative approach, we focused on Egfr signaling, which was recently implicated in ccRCC progression (Liu et al., 2023, Courtney et al., 2018). Intriguingly, while expression of a *UAS-Egfr* transgene in the adult PCs of control males (*C42-Gal4; tub-Gal80ts; UAS-Egfr*) had minimal effects on MT growth and morphology (Figure 10A), expression of this same transgene in the PCs of adult *L2hgdh* mutant males induced significant defects within the lower segment of the MTs, both in terms of tissue organization and ectopic growth (Figure 10B-D). These abnormal cellular structures included small protuberances (Figure 10B), cyst-shaped masses composed of unknown cells attached to MT exterior (Figure 10C,G), and increased cell density within the MT (Figure 10D,G). Moreover, the Egfr-induced ectopic growths readily stained with the lipophilic dye Nile Red (Figure 10 E-G) – an intriguing result considering that human ccRCCs also accumulate elevated amounts of triglycerides. We would also note that the cells within these abnormal masses were of decreased ploidy compared with the PCs and stellate cells (Figure 10B-D,F,G). Considering that we primarily observe these abnormal cell populations in the lower tubule, which contains a population of quiescent renal stem cells (RSCs) that are reactivated in response MT damage (Wang and Spradling, 2020), future studies should examine the possibility that *L2hgdh* mutations alter the proliferation and differentiation of RSCs. Together, these observations demonstrate that loss of L2hgdh activity sensitize the MTs towards experiencing aberrant growth and suggest that this system can be used to evaluate genetic interactions between *L2hgdh* mutations and other genes implicated in both renal cancer and tissue growth.

**Figure 10.**
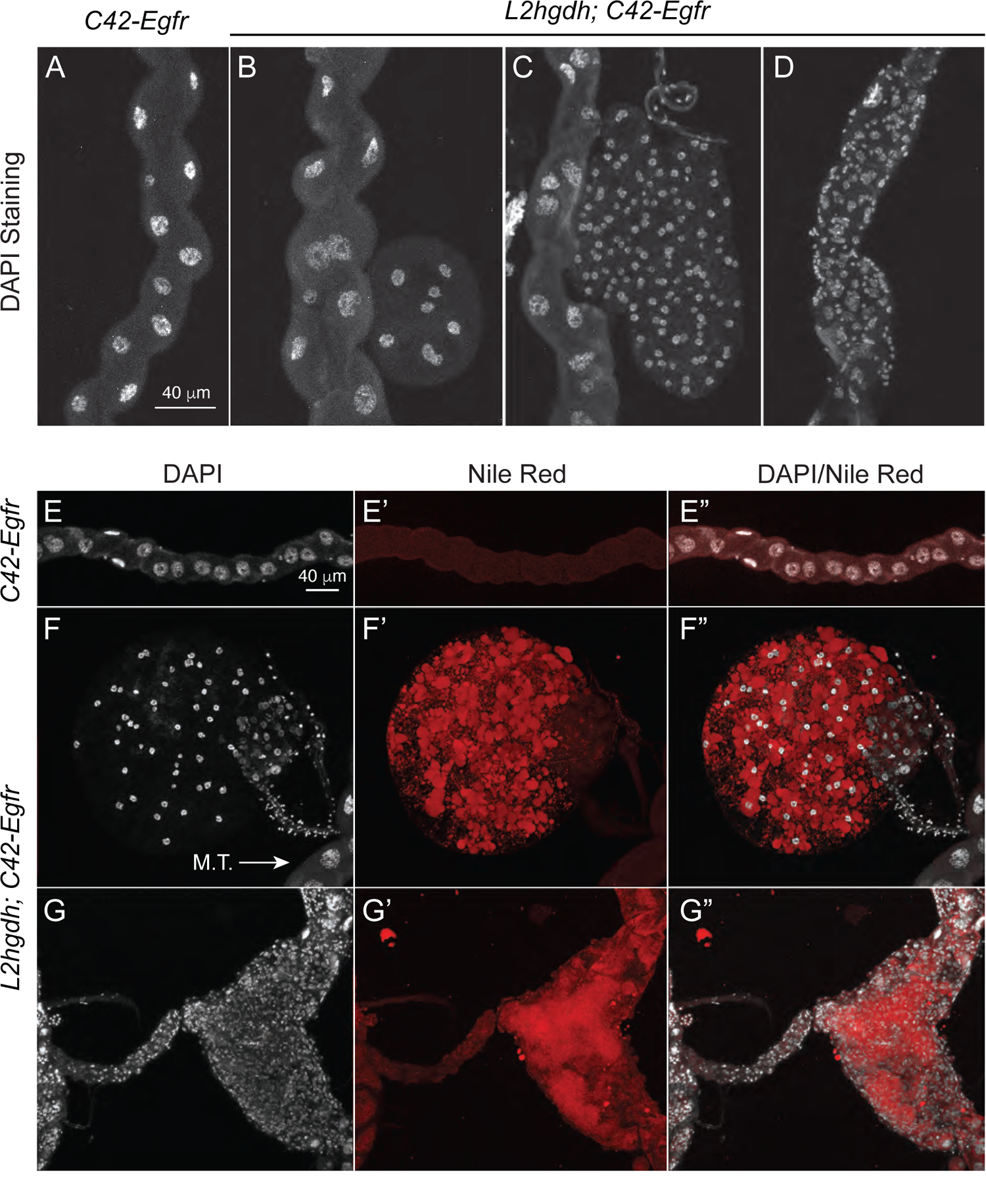
*Egfr* expression induces aberrant tissue growth in *L2hgdh* mutant MTs. Adult male flies were raised at 18°C and shifted to the Gal80ts restrictive temperature (29°C) within 24 hours after eclosion. After 10 days, (A) the lower tubule of *C42-Egfr* adult control male MTs appeared morphologically normal. In contrast, (B-D) *C42-Egfr* expression in a *L2hgdh* mutant background induced (B,C) cysts and (D) abnormal cellular organization. (E-G) MTs expressing *Egfr* in either the control (E) or *L2hgdh* mutant background were stained for (E,F,G) DAPI and (E’, F’, G’) Nile Red to visualize genomic DNA and neutral lipids. Scale bar in (A) applies to (B-D). Scale bar in (E) applies to (F and G).

## DISCUSSION

Here we use the fruit fly *Drosophila melanogaster* to demonstrate that renal L2hgdh activity is sufficient to regulate systemic L-2HG accumulation. In the absence of this enzyme, the resulting elevation in L-2HG levels correlates with increased lactate production and aberrant mitochondrial metabolism. While *L2hgdh* mutants are able to tolerate this metabolic imbalance under standard growth conditions, hypoxia-exposed adult mutant males accumulate far more L-2HG than controls, resulting in a systemic decrease in mitochondrial function and enhanced glycolysis. Moreover, hypoxia-exposed *L2hgdh* mutants are unable to clear the L-2HG buildup and subsequently die during reoxygenation, thus revealing an essential role for L2hgdh in moderating L-2HG levels during hypoxia exposure.

Overall, these results highlight an intimate relationship between L-2HG and oxidative stress. As noted above, animal cells accumulate L-2HG in response to hypoxia, oxidative stress, and mitochondrial dysfunction (Intlekofer et al., 2015, Oldham et al., 2015, Li et al., 2017, Mahmoudzadeh et al., 2020, Hunt et al., 2019, Li and Tennessen, 2019) – findings which suggest that circumstantial L-2HG production might serve a beneficial role. Consistent with these observations, L2hgdh inhibition not only protects cardiac cells against ischemia-reperfusion injury (He et al., 2022), but Dipteran larvae also generate very high L-2HG concentrations during the growth phase (Li et al., 2017). When considered within this context, our findings that *L2hgdh* mutant male adults die following hypoxia exposure seem at odds with a model in which L-2HG benefits the organism. We would note, however, that *L2hgdh* mutants survive hypoxia treatment and specifically die during the reoxygenation phase, suggesting that extremely high L-2HG accumulation in and of itself is not toxic in a low oxygen environment.

When these data are considered in sum with our observation that mitochondrial membrane potential is reduced in *L2hgdh* mutant MTs, a model emerges in which L-2HG accumulation is potentially beneficial. Upon encountering hypoxia, decreased electron transport chain activity results in αKG being shunted into L-2HG production. In turn, L-2HG inhibits αKG dehydrogenase, thus acting as a metabolic failsafe by (i) attenuating mitochondrial metabolism, (ii) shifting cellular metabolism towards a more glycolytic state, and (iii) enhancing oxygen-independent ATP production. Upon return to a normoxic state, however, L-2HG levels must be reduced to properly restore oxidative metabolism. Such a model explains why *L2hgdh* mutant flies die during the reoxygenation phase, as the persistent L-2HG pool would continue to inappropriately suppress oxidative metabolism under aerobic conditions. We would also note that the hypoxia-induced *L2hgdh* mutant phenotypes appear to be independent of altered Hif1α/sima activity, as the very high L-2HG levels present with *L2hgdh* mutants have negligible effects on Hif1α/sima target gene expression.

Our model also provides an explanation for why Dipteran larvae accumulate high levels of L-2HG. Dipteran larvae grow within moist environments that are often hypoxic; however, larvae require aerobic conditions to support growth and development. Based on our current study, we propose that the larval L-2HG pool serves to modulate the swings in mitochondrial metabolism that would inevitably occur as the growing larva moves between hypoxic and normoxic environments. An important test of this model would be to eliminate the larval L-2HG pool and determine if the resulting animals are unable to thrive in their standard food sources.

Our study also raises important questions regarding the role of L2hgdh in renal cancers. Here we demonstrate that the renal system is a key site of L2hgdh activity – not only is renal *L2hgdh* expression sufficient for regulating systemic L-2HG metabolism, but loss of this enzyme within the PCs results in a decrease in mitochondrial membrane potential and elevated ROS production. These results suggest that renal L2hgdh activity restricts organismal L-2HG accumulation – presumably by catalyzing the conversion of L-2HG to αKG. Based on these observations, L-2HG is most likely imported into the PCs for degradation, and in the *L2hgdh* mutant, increased L-2HG levels within these cells directly interferes with PC mitochondrial metabolism.

Moreover, *L2hgdh* mutants respond to elevated L-2HG levels by increasing the MT secretion rate, which likely places increased metabolic demands on an already compromised system. One could imagine how a similar scenario would promote the development and growth of ccRCCs. L-2HG inhibits the TCA cycle in renal cells (Brinkley et al., 2020), and if a cell lacking L2hgdh activity were to experience a sudden bout of hypoxia or oxidative stress, the resulting increase in L-2HG concentrations would shift cellular metabolism from oxidative metabolism towards a more glycolytic state. Considering that ccRCCs are highly glycolytic (Courtney et al., 2018), such a mechanism would further drive the growth and development of these tumors. Our model would also predict that the yet-to-be described L-2HG transporter(s) might also be misregulated in ccRCCs, and future studies should examine this possibility.

In addition to the potential implications for understanding the role of L2hgdh in ccRCCs, our results highlight how environmental conditions could influence the symptoms of individuals experiencing L-2HG aciduria. As noted above, *L2hgdh* mutant flies exhibit few phenotypes when raised under ideal culture conditions. Moreover, the three described *L2hgdh* knockout mouse models are homozygous viable and display a range of metabolic and neurological phenotypes that vary in severity (Rzem et al., 2015, Ma et al., 2017, Brinkley et al., 2020). Such phenotypic variability is also noted in case studies of human L-2HG aciduria patients, where some individuals experience severe neurodevelopmental symptoms in early childhood while other patients experience a milder form of the disease, with disease onset occasionally delayed until middle age.

Our study suggests that environmental stress could, in part, be a cause for this phenotypic variability, as bouts of oxidative stress in L-2HG aciduria patients would induce a hypothetical metabolic feedforward loop where increased L-2HG production inhibits mitochondrial metabolism, which would lead to even more L-2HG production. Considering that L-2HG aciduria is an orphan disease, future studies should examine the role of environmental stress in the etiology of this disease.

One of the most well studied aspects of L-2HG is its role in disrupting the gene expression program (Du and Hu, 2021). Tumors with high levels of L-2HG demonstrate lower levels of genomic 5hmC consistent with TET inhibition (Brinkley et al., 2020, Shim et al., 2014). Similarly, L-2HG can promote Hif1α stability, thus activating gene expression programs that shift the balance of cellular metabolism (Intlekofer et al., 2017, Williams et al., 2022). Our studies suggest that L-2HG has minimal influence on *Drosophila* gene expression – after all, flies exhibit very little DNA methylation, and the three RNAseq experiments described herein failed to identify any gene expression signatures that would be indicative of elevated Hif1α/sima activity. We speculate that the minimal effect of L-2HG on *Drosophila* gene expression is the result of the unique life history of this animal. Since larvae accumulate millimolar concentrations of L-2HG during normal larval development, we hypothesize that *Drosophila* gene regulatory mechanisms have evolved to tolerate high L-2HG levels. Consistent with this hypothesis, Dipteran genomes have lost the gene that encodes DMNT1, and as a result, possess very little DNA methylation (Bewick et al., 2016, Provataris et al., 2018). Thus, the fly provides an ideal model to study how L-2HG influences cellular processes that are often overshadowed by experimental approaches focused on mechanisms that control gene expression.

Finally, our study highlights the ability of L-2HG to influence glycolytic and mitochondrial metabolism, as well as catabolic pathways for lysine and other amino acids that are sensitive to L-2HG abundance. These metabolic pathways are highly conserved between *Drosophila* and humans and the *L2hgdh* mutants provide an ideal genetic platform for studying the interaction between L-2HG and central carbon metabolism. In this regard, our observation that *L2hgdh* mutant MTs accumulate lipids upon *C42-Egfr* expression is of particular interest, as this phenotype mirrors the accumulation of lipids in ccRCCs. These findings, when considered in light of a recent study linking mitochondrial impairment with lipid accumulation in *Drosophila* nephrocytes (the functional equivalent of mammalian proximal tubules)(Lubojemska et al., 2021), establish the fly as a premier system for modeling the metabolic factors that contribute to human renal diseases.

## METHODS

### *Drosophila* husbandry and genetics

Fly stocks were maintained on Bloomington *Drosophila* Stock Center (BDSC) food at 25°C. The strains harboring *L2hgdh^12^* (RRID:BDSC_94700) and *L2hgdh^14^* (RRID:BDSC_94701) were previously described (Li et al., 2017). For our studies, *L2hgdh^14^* was backcrossed six times to a *w^1118^* strain (RRID:BDSC_5905). Unless noted, all experiments were setup by mating 50 virgin *w^1118^; L2hgdh^14^* females with 25 males of either *w^1118^*, *w^1118^; L2hgdh^12^*, or *w^1118^; L2hgdh^12^; p{L2hgdh}* in glass bottles containing BDSC food as previously described (Li and Tennessen, 2017). Male progeny were collected within 8 hrs of eclosion and aged in glass bottles containing BDSC food for 7-10 days at 25°C. Tissue-specific rescue experiments were conducted by crossing *C42-Gal4* (RRID:BDSC_30835) and *elav-Gal4* (RRID:BDSC_8765) into the *L2hgdh^12^* background and crossing males containing these GAL4 drivers with *w^1118^; L2hgdh^14^; UAS-L2hgdh*. The *UAS-L2hgdh* transgene (RRID:BDSC_78053) was previously demonstrated to rescue the *L2hgdh* mutant phenotype (Li et al., 2017). Egfr expression studies were conducted using *UAS-Egfr* (RRID:BDSC_5368) (Zettervall et al., 2004) under the control of *tubP-Gal80^ts^* (RRID:BDSC_7016). Flybase was used throughout this study (Öztürk-Çolak et al., 2024, Gramates et al., 2022).

### Analysis of larval development and pupal mass

White prepupae were individually placed into a pre-tared 1.5 mL microfuge tube and weighed using a Mettler Toledo XS105 balance.

### Climbing Assay

The climbing ability of newly eclosed males was assayed as previously described (Sokol et al., 2008).

### Lactate Dehydrogenase Activity Assays

Ldh activity was measured as previously described with little modification (Rechsteiner, 1970). Briefly, *Drosophila* lysate was prepared from 5-day old flies with the heads removed. Ldh activity was determined at 25°C by monitoring NADH consumption at OD_340_ using a plate reader (BioTek).

### Hypoxia treatment

One day old male flies were put in groups of 20 in vials containing Bloomington *Drosophila* Stock Center standard media, aged for 7-8 days, and incubated in a 1% O_2_ environment as previously described (Mahmoudzadeh et al., 2020). Briefly, the vials were directly placed in a sealed plexiglass chamber that was housed within a 25°C temperature-controlled room. O_2_ concentration within the chamber was controlled and monitored using a ROXY-4 gas regulator (Sable Systems). Nitrogen gas was allowed to flow through a water bubbler and into the chamber until the O_2_ concentration reached 1%. The exhaust valve was then sealed. Relative humidity was maintained at ∼70% by placing 1 L of a saturated NaCl solution (36% NaCl) within the chamber.

### GC-MS analysis

Targeted metabolomics studies were conducted as previously described (Li and Tennessen, 2018, Cox et al., 2017). Briefly, flies were collected in 1.5 mL microfuge tubes and immediately frozen in liquid nitrogen. Samples were transferred to tared 2 mL screwcap tubes containing 1.4 mm ceramic beads that had been pre-chilled in liquid nitrogen. The sample mass was recorded, and tubes were immediately placed back in liquid nitrogen. 800 μL of prechilled (−20 °C) 90% methanol containing 2 µg/mL succinic-d4 acid was added to each sample tube and the sample homogenized in an Omni Beadruptor 24 for 30 seconds at 6.4 m/s. The samples were removed from the homogenizer, incubated at −20°C for 1 hr, and centrifuged at 20,000 x g for 5 min at 4°C. 600 μl of the supernatant was transferred into a new 1.5 mL microcentrifuge tube and dried overnight in a vacuum centrifuge. Dried samples were resuspended in 40 µL of 40 mg/mL methoxylamine hydrochloride (MOX) dissolved in anhydrous pyridine and incubated at 37°C for 1 hour in a thermal mixer shaking at 600 rpm. Samples were then centrifuged for 5 minutes at 20,000 x g and 25 µL of supernatant was transferred into an autosampler vial with a 250 µL deactivated glass microvolume insert (Agilent 5181-8872). 40 µL of N-methyl-N-trimethylsilyltrifluoracetamide (MSTFA) containing 1% TMCS was then added to the sample, at which point the autosampler vial was capped and placed at 37°C for 1 hour with shaking (250 rpm). 1 µL of sample was injected an Agilent GC7890-5977 mass spectrometer equipped with a Gerstel MPS autosampler. Samples were injected with a 10:1 split ratio and an inlet temperature of 300°C. Chromatographic separation was achieved using a 0.25 mm x 30 m Agilent HP-5ms Ultra Insert GC column with a helium carrier gas flow rate of 1.98 mL/min. The GC temperature gradient was as follows: (1) Hold at 95°C for 1 min. (2) Increase temperature to 110°C with a 40°C/min ramp. Hold 2 min. (3) Increase temperature to 250°C with a 25°C/min ramp. (4) Increase temperature to 330°C with a 25°C/min ramp and hold for 4 minutes.

### D-/L-2HG Quantification

Sample collection, homogenization, and 2HG quantification were conducted as previously described (Li and Tennessen, 2019). Briefly, 800 μL of extraction buffer (90% MeOH containing 8 mg of 2,2,3-d3-R,S-2-hydroxyglutarate) was added to each sample tube (kept in −20°C enzyme caddies). Tubes were then placed in a Omni Bead Ruptor 24 and the sample homogenized for 30 seconds at 6.4 m/s. Samples were subsequently incubated at −20°C for 1 hr. After the incubation, samples were centrifuged at 20,000 x g for 5 min at 4°C. 600 μl of the supernatant was then transferred into a 1.5 mL microfuge tube and dried overnight using a vacuum centrifuge.

Dried samples were resuspended in 50 μl of R-2-Butanol and 5 μl of HCL and incubated at 90°C for three hours with shaking at 300 rpm using an Eppendorf ThermoMixer F1.5. After cooling, 200 μl of water and 500 μl of hexane was added to each tube. The organic phase (hexane) was then transferred to a new tube and dried for 30 minutes using a vacuum centrifuge. The dried samples were resuspended in 60 μl of acetic anhydride and 60 μl of pyridine and incubated at 80°C for 1 hr with shaking at 300 rpm. Samples were then dried for 3 hrs using a vacuum centrifuge and resuspended in 60 μl of hexane.

Derivatized samples were analyzed using a 7890B-5977B MSD Agilent GC-MS. 1 μl of the sample was injected into a 30m DB-5MS column with a 0.25 mm inner diameter. The GC temperature gradient was as follows: (1) Initial oven temperature was set to 95°C with a 1 min hold time. (2) Increased temperature to 110°C at a rate of 40°C/min. (3) Increase temperature to 200°C at a rate of 5°C/min. (4) Increase temperature to 330°C at a rate of 40°C/min. The GC-MS was set to Selected Ion Monitoring (SIM) dwelling on 173 and 176 m/z value. MassHunter Quantitative software was used to measure the amount of D-/L-2HG present within each sample.

### Ultra High-pressure Liquid Chromatography - Mass Spectrometry (UHPLC-MS)-based Metabolomics

UHPLC-MS metabolomics analyses were performed at the University of Colorado Anschutz Medical Campus, as previously described (Nemkov et al., 2019). Briefly, the analytical platform employs a Vanquish UHPLC system (Thermo Fisher Scientific, San Jose, CA, USA) coupled online to a Q Exactive mass spectrometer (Thermo Fisher Scientific, San Jose, CA, USA). The (semi)polar extracts were resolved over a Kinetex C18 column, 2.1 x 150 mm, 1.7 µm particle size (Phenomenex, Torrance, CA, USA) equipped with a guard column (SecurityGuard^TM^ Ultracartridge – UHPLC C18 for 2.1 mm ID Columns – AJO-8782 – Phenomenex, Torrance, CA, USA) using an aqueous phase (A) of water and 0.1% formic acid and a mobile phase (B) of acetonitrile and 0.1% formic acid for positive ion polarity mode, and an aqueous phase (A) of water:acetonitrile (95:5) with 1 mM ammonium acetate and a mobile phase (B) of acetonitrile:water (95:5) with 1 mM ammonium acetate for negative ion polarity mode. The Q Exactive mass spectrometer (Thermo Fisher Scientific, San Jose, CA, USA) was operated independently in positive or negative ion mode, scanning in Full MS mode (2 μscans) from 60 to 900 m/z at 70,000 resolution, with 4 kV spray voltage, 45 sheath gas, 15 auxiliary gas. Calibration was performed prior to analysis using the Pierce^TM^ Positive and Negative Ion Calibration Solutions (Thermo Fisher Scientific).

### Statistical Analysis of Metabolomics Data

All metabolomics datasets were analyzed using Metaboanalyst 5.0 (Pang et al., 2021), with data normalized to sample mass and preprocessed using log normalization and Pareto scaling. Data generated by GC-MS analysis of individual compounds was analyzed using GraphPad Prism 10.

### Collection of *Drosophila* Excreta for GC-MS analysis

Male flies were collected within one day of eclosion, transferred into vials with Bloomington food, and aged for 7 days. Groups of 10 male flies were then placed into a 1.5 mL microfuge tube with a hole poked in the lid to allow gas exchange. Tubes were put in an incubator (25°C, relative humidity ∼80%) for 24 hours. After the 24 hr incubation period, flies were discarded and the sides of the tubes were rinsed using chilled extraction buffer. The extraction buffer was then processed for D/L-2HG quantification.

### RNA-seq data analysis

For all genotypes, RNA-seq was performed on 3 whole-fly biological replicates for each condition/genotype, with each sample containing either 20 adult flies or 100 dissected MTs. Samples were paired-end sequenced on the NextSeq 75 platform to a depth of 15-20 million reads each, using standard Illumina TruSeq Stranded mRNA libraries, at the IU Center for Bioinformatics and Genomics (CGB).

### L2hgdh whole body and Malpighian Tubule datasets

Analysis was performed as described previously using a python based in-house pipeline (https://github.com/jkkbuddika/RNA-Seq-Data-Analyzer) (Buddika et al., 2021). First, the quality of raw sequencing files was assessed using FastQC version 0.11.9 (Andrews, 2010), and reads with low quality were removed using Cutadapt version 2.9 (Martin, 2011). Subsequently, the remaining high-quality reads were aligned to the Berkeley *Drosophila* Genome Project (BDGP) assembly release 6.32 (Ensembl release 103) reference genome using STAR genome aligner version 2.7.3a (Dobin et al., 2013).

Additionally, duplicated reads were eliminated using SAMtools (Li et al., 2009) version 1.10. Finally, the Subread version 2.0.0 (Liao et al., 2019), function *featureCounts*, was used to count the number of aligned reads to the nearest overlapping feature. All subsequent downstream analyses and data visualization steps were performed using custom scripts written in R. To identify differentially expressed genes in different genetic backgrounds the Bioconductor package DESeq2 (https://bioconductor.org/packages/release/bioc/html/DESeq2.html) version 1.26.0 was used (Love et al., 2014). Unless otherwise noted, significantly upregulated and downregulated genes were defined as FDR < 0.05; Log_2_ fold change > 1 and FDR < 0.05; Log_2_ fold change < −1.

### Hypoxia timecourse experiments

RNA-seq read quality was assessed with FastQC (Andrews, 2010) and MultiQC (Ewels et al., 2016), with adequate reads and no significant issues noted. Reads were pseudo-aligned and quantified using Kallisto v0.46.0 (Bray et al., 2016), the *D. melanogaster* BDGP6.32 reference assembly and annotation retrieved through Ensembl (Cunningham et al., 2022), and the gffread utility from the Cufflinks suite (Trapnell et al., 2010) to generate the transcriptome from the assembly and annotation.

Differential expression analysis was performed with DESeq2 v1.30.1 (Love et al., 2014) running in RStudio v1.4.1717 on R v4.0.4. Genes with average expression below 2 counts per sample (i.e. 48 counts per row) were filtered out before determining genes with significant expression difference in at least one sample with the likelihood-ratio test (LRT) and cutting off at LRT *p* ≤ 0.05. All subsequent experiment-wide analyses were applied to this set of normalized, significant gene counts, which were ranked by variance and further cutoff as noted in the figures.

Subsequent analysis and visualization were performed in R using a variety of tools and packages. Correlation plots were generated using base R and the corrplot package (Wei and Simko, 2021). Principal components were analyzed and visualized with PCAtools v2.2.0 (Blighe and Lun, 2022). Heatmaps, including clustering analysis performed by the hclust method (from base R stats package), were generated by pheatmap (Kolde, 2019).

### PANGEA Analysis

All RNAseq data were analyzed using PANGEA (Hu et al., 2023). Genes that were significantly down- or up-regulated were analyzed for Gene Ontology Enrichment using the SLIM2 GO BP and FlyBase signaling pathway (experimental evidence) sets (Consortium et al., 2023).

### Ramsay Assay

The following genotypes were used: *w^1118^; L2hgdh^14/+^* (control), *w^1118^; L2hgdh^12/14^*(experimental), and *w^1118^; L2hgdh^12/14^; p{L2hgdh}* (rescue). Mated adult females were collected and maintained at 25°C and fed with standard fly food prepared in a central kitchen at the University of Utah and supplemented with yeast. Flies were transferred to fresh food every 2-4 days. Anterior Malpighian tubules were isolated on days 3-9 post-eclosion. Fluid secretion over 2 hours in standard bathing medium was measured using the Ramsay assay as described previously (Schellinger and Rodan, 2015). Standard bathing medium consists of a 1:1 mixture of Schneider’s medium (Gibco, 21720024) and *Drosophila* saline (composition in mM: NaCl 117.5, KCl 20, CaCl_2_ 2, MgCl_2_ 8.5, NaHCO_3_ 10.2, NaH_2_PO_4_ 4.3, HEPES 15, and glucose 20, pH 7.0). Statistical testing was performed using GraphPad Prism 10.

### JC-1 Staining

Adult Malpighian tubules were dissected in Schneider’s medium. Immediately following dissection, JC-1 was added to the media at a 1:1000 dilution and Malpighian tubules were incubated for 10 minutes. Samples were then washed twice with Schneider’s medium, mounted immediately in VECTASHIELD, and visualized using the Leica SP8 confocal microscope housed within the Indiana Light Microscopy Facility.

### ROS (DHE) Staining

Flies were collected and Malpighian tubules were dissected in Schneider’s medium. DHE 30mM stock solution was freshly made. Dissected samples were incubated in Schneider’s medium containing 60uM DHE for 5 minutes. Samples were washed three times in Schneider’s medium for 5 minutes and were mounted immediately in vectashield. Confocal microscopy was performed immediately.

### *Egfr* expression and Nile Red Staining

Malpighian tubules of adult flies were dissected in 1X PBS and fixed in 4% formaldehyde in 1X PBS for 30 minutes. Tissues were washed with 1X PBS for 10 minutes after which they were incubated in 1:2000 dilution of 0.5 mg/ml Nile Red (Sigma Aldrich-N3013) with PBS for 30 minutes. Subsequently, tissues were rinsed three times with 1X PBS and mounted in Vectashield mounting medium with DAPI (Vector Labs) for nuclei staining. Samples were examined under SP8 fluorescence microscope.

### Renal Cell Culture Experiments

786-O and OS-RC-2 cells were maintained in RPMI media (Corning #10041CV) supplemented with 10% FBS (R&D system #S11150) and 1X penicillin-streptomycin (Gibco #15140122). Cells were periodically tested and confirmed to be free of mycoplasma contamination. Lentiviral particles harboring control vector or L2HGDH construct were used to transduce the cells and selected by puromycin (1 ug/ ml) as described (Shelar et al., 2018). The selected polyclonal pools were verified for L2HGDH overexpression by immunoblot. ∼1.5 million control/L2HGDH cells (n=6, each) were cultured for 24 hrs. After 24 hrs, the spent media was aspirated out after aliquoting and rapidly freezing (in LN2) 1 ml media from each condition. The aliquoted spent media were stored in −80 deg C until used for the L-2HG assays.

## ACKNOWLEDGEMENTS

Thanks to the Bloomington *Drosophila* Stock Center (NIH P40OD018537) for providing fly stocks, the *Drosophila* Genomics Resource Center (NIH 2P40OD010949) for genomic reagents, and Flybase (NIH 5U41HG000739). Metabolomics analysis performed at the Metabolomics Core Facility at the University of Utah is supported by spectrometry equipment obtained through NCRR Shared Instrumentation Grants 1OD016232-01, 1S10OD018210-01A1 and 1S10OD021505-01. Sequencing was performed at Indiana University’s Center for Genomics and Bioinformatics. All computation was performed on Indiana University’s Carbonate and Research Desktop HPC clusters. S.S. is supported by NCI R01CA200653 and NCI F30CA232397. A.K. was supported in part by a Research Scholar Award from the Urology Care Foundation (Award No.669279) and a concept award from the Department of Defense (Award No. HT9425-23-1-0339). GC acknowledges support from the DBT/Wellcome Trust India Alliance Fellowship/Grant [grant number IA/I(S)/17/1/503085]. A.R.R. is supported by the National Institute of Diabetes and Digestive and Kidney Diseases (R01DK110358). J.M.T. is supported by the National Institute of General Medical Sciences of the National Institutes of Health under a R35 Maximizing Investigators’ Research Award (MIRA; 1R35GM119557).

## SUPPLEMENTAL FIGURE LEGENDS

**Figure S1. *L2hgdh* mutant larvae and pupae grow at the same rate as controls strains.** *L2hgdh^14/+^* controls, *L2hgdh^12/14^* mutants, and *L2hgdh^12/14^; p{L2hgdh}* rescued animals exhibit no significant differences in (A) the duration of larval development or the body mass of (B) white prepupae, (C) newly eclosed males, or (D) newly eclosed females.

**Figure S2. *L2hgdh* mutant adult males exhibit normal climbing behavior and lifespan.** (A) *L2hgdh^14/+^* control, *L2hgdh^12/14^* mutant, and *L2hgdh^12/14^; p{L2hgdh}* rescued adult males exhibit similar climbing abilities. (B) Adult male *w^1118^* controls and genetically-matched *L2hgdh^14^* mutants show no difference in lifespan.

**Figure S3. RNAseq analysis of *L2hgdh* mutant adult males.** RNA isolated from *L2hgdh^14/+^* control and *L2hgdh^12/14^* mutant adult males was analyzed using RNAs eq. 2 40 genes were significantly down-regulated and 119 genes were up-regulated. (A) Of these significantly altered genes, 35 metabolic genes were down-regulated and 9 metabolic genes were up-regulated. In addition, mutant males exhibited no significant changes in the expression of genes involved in (A) glycolysis or (B) the TCA cycle. Only one gene involved in (D) the electron transport chain (ETC) was significantly mis-regulated in *L2hgdh^12/14^* mutants.

**Figure S4.** A comparison of the metabolomic data from *L2hgdh^14/+^* control and *L2hgdh^12/14^* mutant samples using principal component analysis (PCA). Metaboanalyst was used to analyze the metabolomics data from *L2hgdh^14/+^* control and *L2hgdh^12/14^* mutant adult male samples using PCA.

**Figure S5. Targeted GC-MS analysis of *L2hgdh* mutants.** GC-MS analysis of *L2hgdh^14/+^* control, *L2hgdh^12/14^* mutant, and *L2hgdh^12/14^; p{L2hgdh}* rescued adult males revealed that (A) 2HG, (B) lactate, and (D) lysine levels were elevated in mutant samples. However, (C) pyruvate levels were unchanged in these experiments. Similarly, GC-MS analysis of hemolymph extracted adult males of the same genotypes contained elevated levels of (E) 2HG and (F) lactate (Lac). Data presented as a scatter plot with the center line representing the mean and error bars representing the standard deviation. *\*\*\*P*<0.001. Statistical analysis conducted using an ordinary one-way ANOVA followed by a Dunnett’s multiple comparison test.

**Figure S6. *L2hgdh* mutant exhibit normal levels of LDH activity.** Lactate dehydrogenase (LDH) activity was assessed in whole animal extracts of *L2hgdh^14/+^* controls, *L2hgdh^12/14^* mutants, and *L2hgdh^12/14^; p{L2hgdh}* rescued animals. (A,B) Extracts from both control and mutant animals possess similar levels of LDH activity. However, LDH activity in extracts from rescued animals were significantly altered. (A) Extracts from rescued animals converted pyruvate-to-lactate at a lower rate than extracts from either the heterozygous control or *L2hgdh* mutant strain. (B) Extracts from rescued animals converted lactate-to-pyruvate at a higher rate than extracts from either the heterozygous control or *L2hgdh* mutant strain. Histograms represent the mean and error bars represent standard deviation. *\*P*<0.01, *\*\*\*P*<0.001.

**Figure S7. Hierarchical clustering analysis reveals that *L2hgdh* mutant metabolome is significantly disrupted by hypoxia exposure.** A Hierarchical Clustering Dendrogram of the metabolomics samples from Table S4. All normoxia samples, regardless of genotype cluster in a single clade. Following hypoxia exposure, however, the control and rescue samples cluster into distinct clades – one representing the H+0 timepoint and the second representing the H+1 timepoint. In contrast, the H+0 and H+1 mutant samples cluster together, thus illustrating the similarities between the mutant metabolomes at these two timepoints. MetaboAnalyst 5.0 was used to analyze the data and the dendrogram was generated using Euclidean Distance and Ward’s method for the Clustering Algorithm.

**Figure S8** Metabolites associated with glycolysis are elevated in *L2hgdh* mutants following hypoxia exposure. The relative abundance of (A) 2-hydroxyglutarate (2HG), (B) glucose, (C) pyruvate and (D) lactate in *L2hgdh^12/14^* mutants, and *L2hgdh^12/14^; p{L2hgdh}* rescued adult male flies following either (i) a 12 hr incubation under normoxic conditions, (ii) a 12 hr incubation in 1% O_2_ (H), or (iii) a 12 hr incubation in 1% O_2_ followed by a 1 hr recovery in normoxic conditions (H+1 hr). While only (A) 2HG levels were elevated in *L2hgdh^12/14^* mutants under normoxia relative to the heterozygous control and *L2hgdh^12/14^; p{L2hgdh}* rescue line, all four metabolites (A-D) were significantly elevated in mutant samples at the H+0 and H+1 timepoint. Data normalization and statistical analysis conducted using an ANOVA followed by a Fisher’s least significant difference test in MetaboAnalyst 5.0.

**Figure S9. *L2hgdh* mutants exhibit significant decreases in TCA cycle metabolites following hypoxia exposure.** The relative abundance of (A) citrate, (B) succinate, (C) fumarate and (D) malate in *L2hgdh^12/14^* mutants, and *L2hgdh^12/14^; p{L2hgdh}* rescued adult male flies following either (i) a 12 hr incubation under normoxic conditions, (ii) a 12 hr incubation in 1% O_2_ (H), or (iii) a 12 hr incubation in 1% O_2_ followed by a 1 hr recovery in normoxic conditions (H+1 hr). Under normoxic conditions, all four metabolites were present at similar levels in *L2hgdh^12/14^* mutants, heterozygous controls, and *L2hgdh^12/14^; p{L2hgdh}* rescue animals. In contrast, all four metabolites were present at significantly lower levels in *L2hgdh^12/14^* mutants at the H+0 and H+1 timepoints when compared with the control and rescue samples. Data normalization and statistical analysis conducted using an ANOVA followed by a Fisher’s least significant difference test in MetaboAnalyst 5.0.

**Figure S10.** *L2hgdh* mutants exhibit significant changes in amino acid levels following hypoxia exposure. The relative abundance of (A) glutamate, (B) aspartate, (C) proline and (D) serine in *L2hgdh^12/14^* mutants, and *L2hgdh^12/14^; p{L2hgdh}* rescued adult male flies following either (i) a 12 hr incubation under normoxic conditions, (ii) a 12 hr incubation in 1% O_2_ (H), or (iii) a 12 hr incubation in 1% O_2_ followed by a 1 hr recovery in normoxic conditions (H+1 hr). Under normoxic conditions, all four amino acids were present at similar levels in *L2hgdh^12/14^* mutants, heterozygous controls, and *L2hgdh^12/14^; p{L2hgdh}* rescue animals. In contrast, all four metabolites were present at significantly lower levels in *L2hgdh^12/14^* mutants at the H+0 and H+1 timepoints when compared with the control and rescue samples. In addition, (D) serine levels were elevated at the H+0 and H+1 timepoint for both the control and rescue strain relative when compared with normoxic samples, but decreased in mutant samples at the H+0 and H+1 timepoints. Data normalization and statistical analysis conducted using an ANOVA followed by a Fisher’s least significant difference test in MetaboAnalyst 5.0.

**Figure S11. Additional analysis of RNAseq data from hypoxia exposed *L2hgdh* mutants.** (A) Eigencor plot of PC correlations with experimental variables (Kendall’s tau). Asterisks after correlation value denote statistical significance. (B-D) Expression levels of *Hph*, *Ldh*, and *Thor* for each genotype across the timecourse. Values are mean linearized expression derived from the normalized, transformed log counts of each set of replicates. Error bars represent standard error of the mean. Each gene presented has LRT adjusted p-value << 0.05.

**Figure S12. *L2hgdh* expression in the nervous system does not prevent *L2hgdh* mutants from dying during reoxygenation.** Adult male flies were exposed to 1% O_2_ for 12 hours and monitored for recovery over the course of 90 minutes. While nearly all *L2hgdh^14/+^* control flies recovered from hypoxia treatment, over 75% of *L2hgdh^12/14^* mutant failed to recover. *L2hgdh^12/14^; elav-Gal4 +/+ UAS-L2hgdh*, which express *L2hgdh* within the nervous system, exhibit no increase in survival when compared with the negative control genotypes *L2hgdh^12/14^; elav-Gal4/+* and *L2hgdh^12/14^; UAS-L2hgdh/+*. n=6 vials containing 10 adult male flies for each genotype.

**Figure S13. L-2HG levels are elevated in the urine of *L2hgdh* mutant mice.** L-2HG levels were quantified in urine from control and *L2hgdh* mutant mice, as described in Brinkley, et al., 2020.

**Figure S14. RNAseq analysis of MTs from *L2hgdh* mutant males.** RNA isolated from the MTs of *L2hgdh^14/+^* control and *L2hgdh^12/14^* mutant adult males was analyzed using RNAseq. (A) A volcano plot illustrating comparing RNA abundance in *L2hgdh^12/14^*mutants relative to *L2hgdh^14/+^* controls. 90 genes were significantly down-regulated and 124 genes were up-regulated. (B-C) Only one gene involved in (B) glycolysis, (C) the TCA cycle, or (D) the electron transport chain (ETC) was significantly mis-regulated in *L2hgdh^12/14^* mutants.

## Supplemental Tables

**Table S1.** RNA-seq results comparing gene expression between adult male *L2hgdh^12/14^*mutants and *L2hgdh^14/+^* controls. All genes are included in this table, regardless of whether the change in gene expression was significant.

**Table S2.** RNA-seq results comparing gene expression between adult male *L2hgdh^12/14^*mutants and *L2hgdh^14/+^* controls. Only genes displaying significant changes in gene express are included in this table.

**Table S3.** A list of metabolic genes that were significantly up- or downregulated in Table S2.

**Table S4.** PANGEA analysis of significantly misregulated genes in *L2hgdh^12/14^* mutants as compared with *L2hgdh^14/+^* controls (see Table S3 for list of genes used in the analysis. PANGEA default settings (FlyBase signaling pathway; SLIM2 GO BP) were used to analyze for enrichment. Only Gene Set IDs with a *P-*value of less than 0.1 are included in the table.

**Table S5.** Metabolomic analysis of *L2hgdh^12/14^* mutants as compared with *L2hgdh^14/+^*controls. All values normalized to sample mass and a d4-succinic acid standard. n=6 samples per genotype; 20 adult males per sample.

**Table S6.** Metabolomic Analysis of L2hgdh mutants following hypoxia treatment. *L2hgdh^14/+^* controls, *L2hgdh^12/14^* mutants, and *L2hgdh^12/14^; p{L2hgdh}* rescued adult males were exposed to either normoxic or hypoxic conditions and collected at the following three timepoints: (i) 12 hr incubation in normoxic conditions (Norm), (ii) immediately following a 12 hr hypoxia exposure (H+0 hr), and (iii) following a 1 hr recovery period in the presence of atmospheric oxygen (H+1 hr). All samples contained 20 adult males. Data are normalized to sample mass and an internal d4-succinic acid standard.

**Table S7.** RNAseq analysis of *L2hgdh* mutants following hypoxia treatment. *L2hgdh^14/+^* controls and *L2hgdh^12/14^* mutants adult males were exposed to either normoxic or hypoxic conditions and collected at the following four timepoints: (i) 12 hr incubation in normoxic conditions (Norm), (ii) immediately following a 12 hr hypoxia exposure (H+0 hr), (iii) following a 1 hr recovery period in the presence of atmospheric oxygen (H+1 hr), and (iv) following a 12 hr recovery period in the presence of atmospheric oxygen (H+12 hr). All genes are included in this table, regardless of whether the change in gene expression was significant.

**Table S8.** RNAseq analysis of *L2hgdh^12/14^* mutants following hypoxia treatment. *L2hgdh^14/+^* controls and *L2hgdh^12/14^* mutants adult males were exposed to either normoxic or hypoxic conditions and collected at the following four timepoints: (i) 12 hr incubation in normoxic conditions (Norm), (ii) immediately following a 12 hr hypoxia exposure (H+0 hr), (iii) following a 1 hr recovery period in the presence of atmospheric oxygen (H+1 hr), and (iv) following a 12 hr recovery period in the presence of atmospheric oxygen (H+12 hr). Only genes displaying significant changes in gene express are included in this table.

**Table S9.** RNAseq analysis of *L2hgdh^14/+^* controls exposed to either normoxic or hypoxic conditions and collected at the following four timepoints: (i) 12 hr incubation in normoxic conditions (Norm), (ii) immediately following a 12 hr hypoxia exposure (H+0 hr), (iii) following a 1 hr recovery period in the presence of atmospheric oxygen (H+1 hr), and (iv) following a 12 hr recovery period in the presence of atmospheric oxygen (H+12 hr). All genes are included in this table, regardless of whether the change in gene expression was significant.

**Table S10.** RNAseq analysis of *L2hgdh^14/+^* controls exposed to either normoxic or hypoxic conditions and collected at the following four timepoints: (i) 12 hr incubation in normoxic conditions (Norm), (ii) immediately following a 12 hr hypoxia exposure (H+0 hr), (iii) following a 1 hr recovery period in the presence of atmospheric oxygen (H+1 hr), and (iv) following a 12 hr recovery period in the presence of atmospheric oxygen (H+12 hr). Only genes displaying significant changes in gene express are included in this table.

**Table S11.** RNAseq analysis of *L2hgdh^12/14^* mutants exposed to either normoxic or hypoxic conditions and collected at the following four timepoints: (i) 12 hr incubation in normoxic conditions (Norm), (ii) immediately following a 12 hr hypoxia exposure (H+0 hr), (iii) following a 1 hr recovery period in the presence of atmospheric oxygen (H+1 hr), and (iv) following a 12 hr recovery period in the presence of atmospheric oxygen (H+12 hr). All genes are included in this table, regardless of whether the change in gene expression was significant.

**Table S12.** RNAseq analysis of *L2hgdh^12/14^* mutants exposed to either normoxic or hypoxic conditions and collected at the following four timepoints: (i) 12 hr incubation in normoxic conditions (Norm), (ii) immediately following a 12 hr hypoxia exposure (H+0 hr), (iii) following a 1 hr recovery period in the presence of atmospheric oxygen (H+1 hr), and (iv) following a 12 hr recovery period in the presence of atmospheric oxygen (H+12 hr). Only genes displaying significant changes in gene express are included in this table.

**Table S13.** PANGEA analysis of significantly misregulated genes in *L2hgdh^12/14^* mutants as compared with *L2hgdh^14/+^* controls. Experiment 1 = Norm_Mutant_vs_Control; Experiment 2 = H+0_Mutant_vs_Control; Experiment 3 = H+1_Mutant_vs_Control; Experiment 4 = H+12_Mutant_vs_Control;

**Table S14.** RNA-seq results comparing Malpighian tubule gene expression between adult male *L2hgdh^12/14^* mutants and *L2hgdh^14/+^* controls. All genes are included in this table, regardless of whether the change in gene expression was significant.

**Table S15.** RNA-seq results comparing Malpighian tubule gene expression between adult male *L2hgdh^12/14^* mutants and *L2hgdh^14/+^* controls. Only genes displaying significant changes in gene express are included in this table.

**Table S16.** PANGEA analysis of significantly misregulated genes in the MTs of *L2hgdh^12/14^* mutants as compared with *L2hgdh^14/+^* controls. (see Table S15 for list of genes used in the analysis. PANGEA default settings (FlyBase signaling pathway; SLIM2 GO BP) were used to analyze for enrichment. Only Gene Set IDs with a *P-*value of less than 0.1 are included in the table.

