## Supplemental Figures for "Renal L-2-hydroxyglutarate dehydrogenase activity promotes hypoxia tolerance and mitochondrial metabolism in *Drosophila melanogaster*"

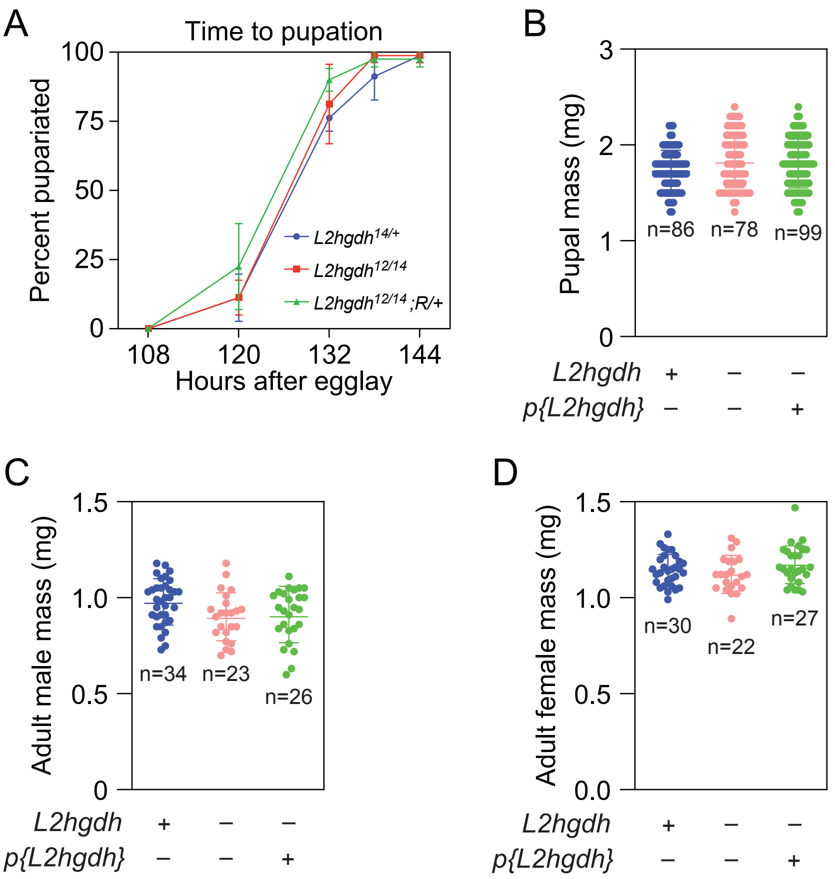


Supplemental Figure 1

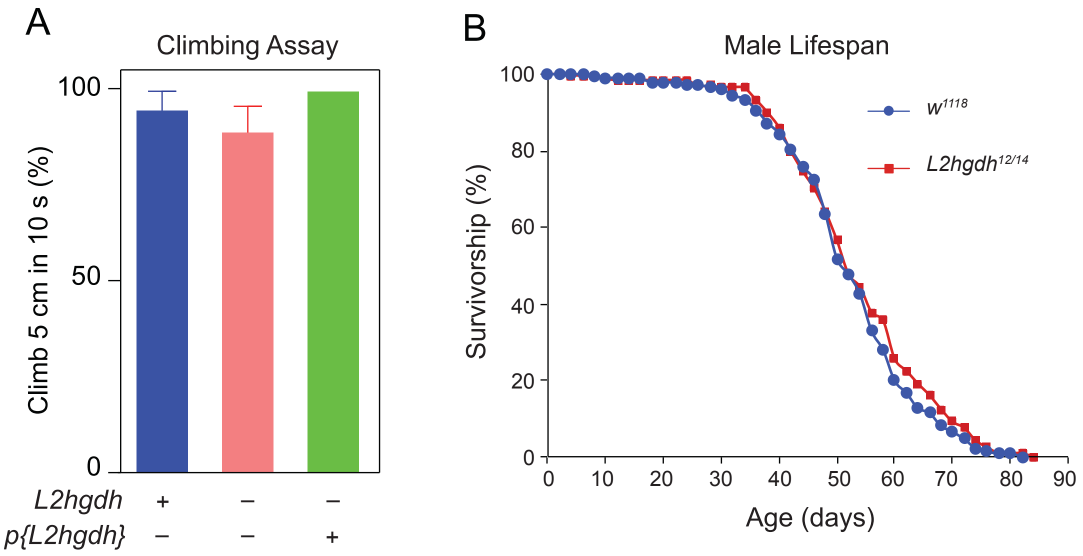


Supplemental Figure 2

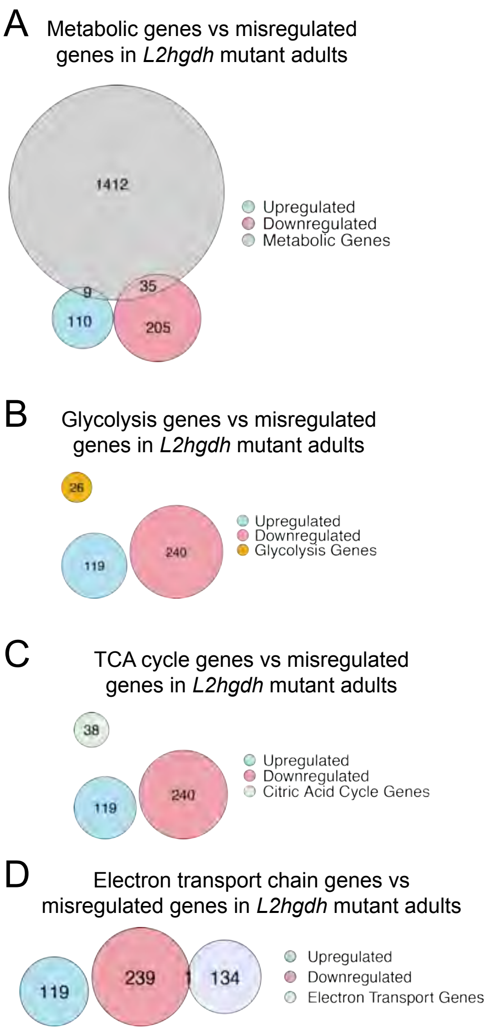


Supplemental Figure 3

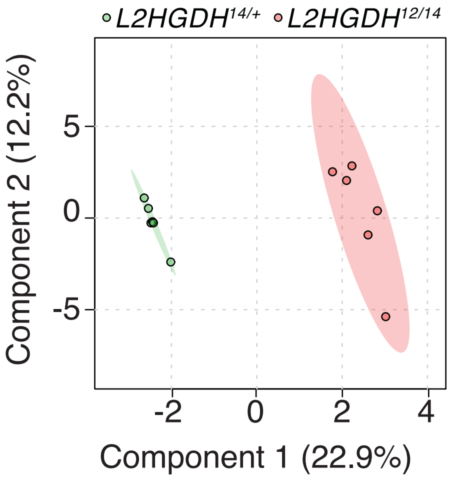


Supplemental Figure 4

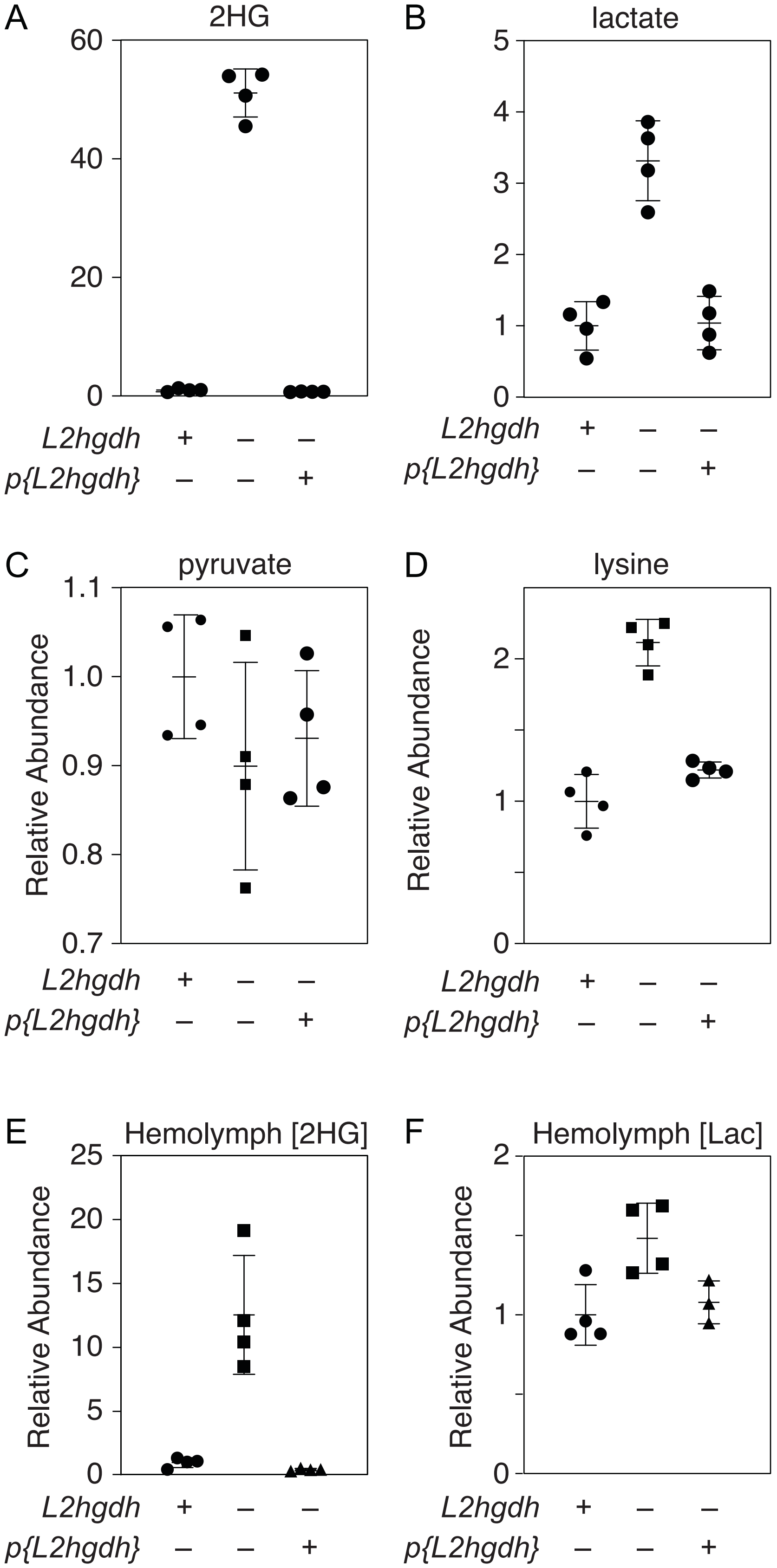


Supplemental Figure 5

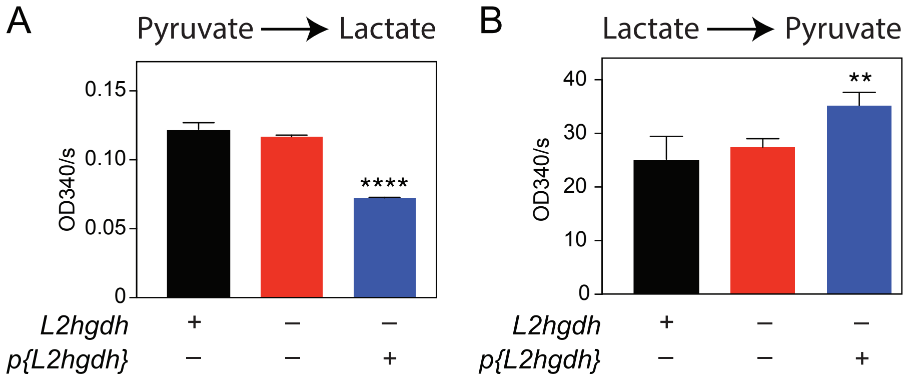


Supplemental Figure 6

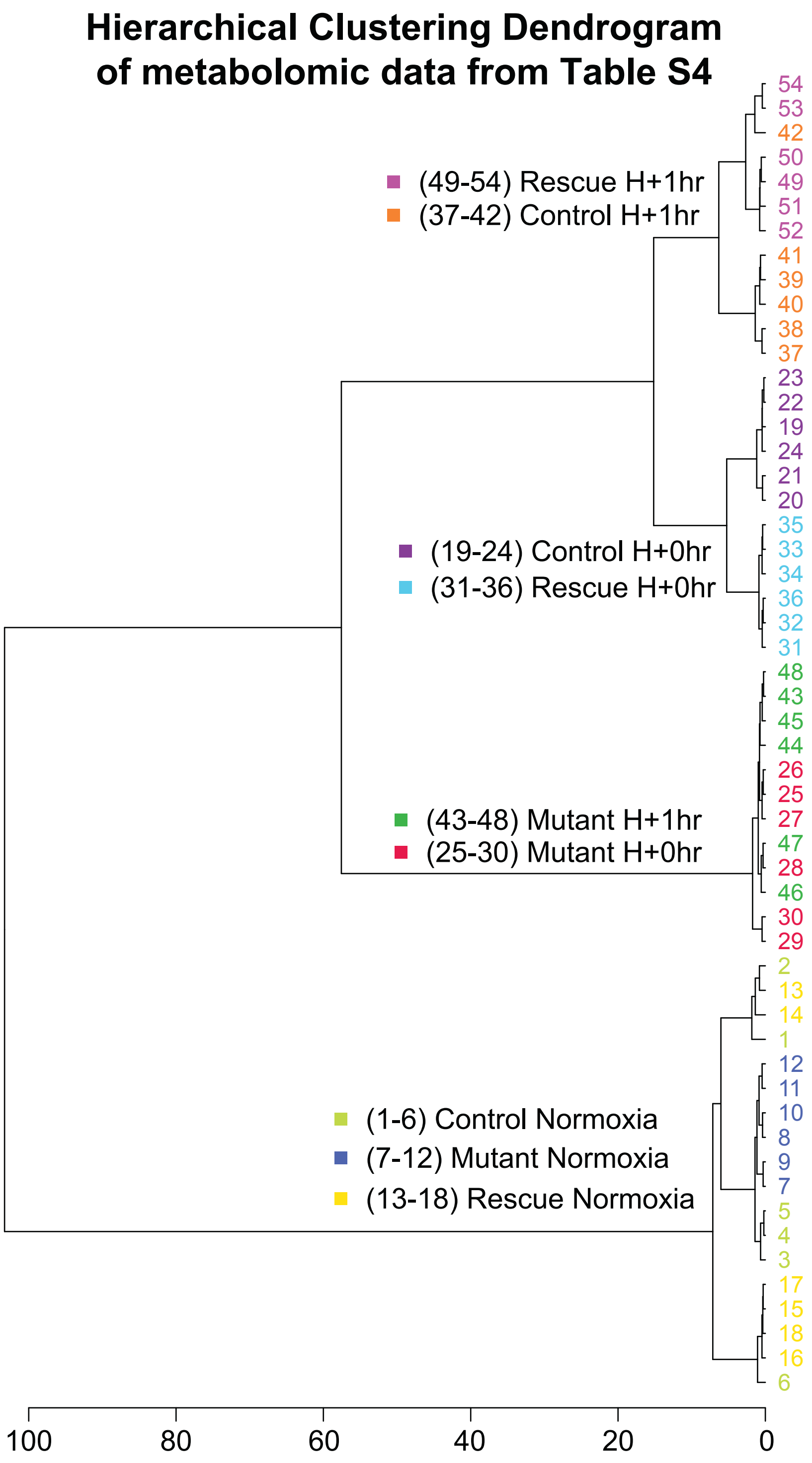


Supplemental Figure 7

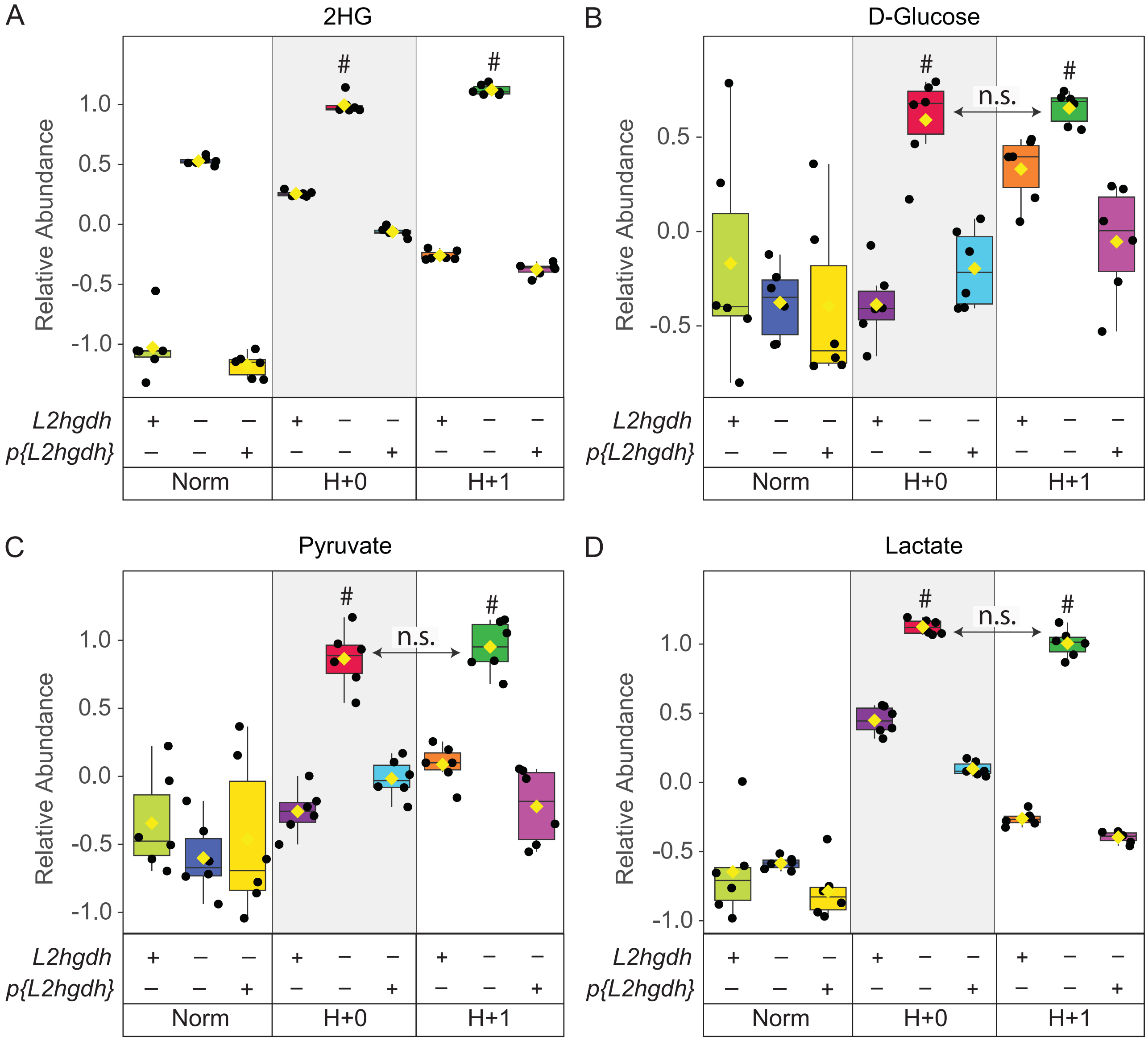

### - Unless noted, *P*>0.05 when compared with all other genotypes and conditions.

Supplemental Figure 8

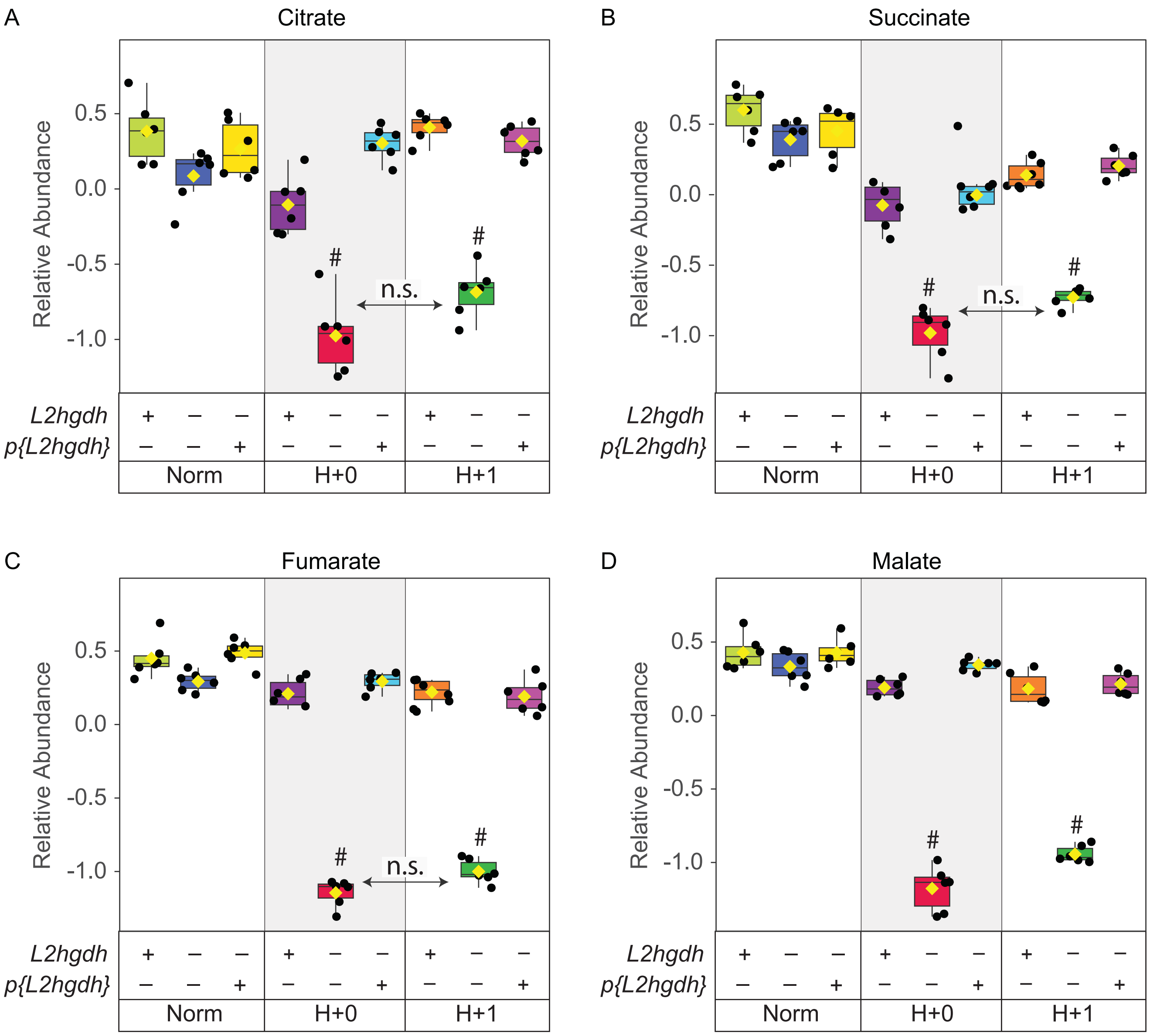

### - Unless noted, $P$<0.05 when compared with all other genotypes and conditions.

Supplemental Figure 9

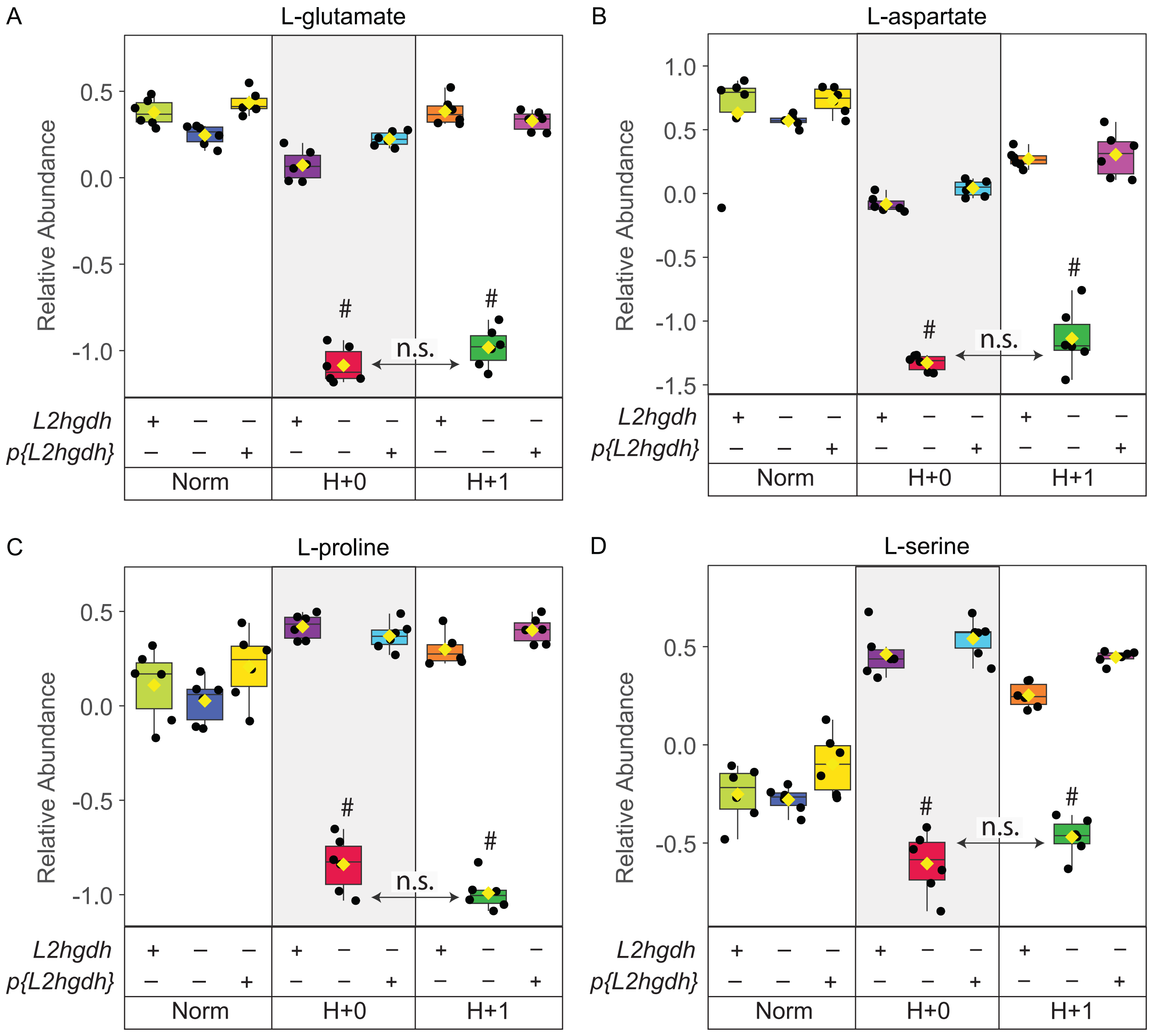


### - Unless noted, *P*<0.05 when compared with all other genotypes and conditions.

Supplemental Figure 10

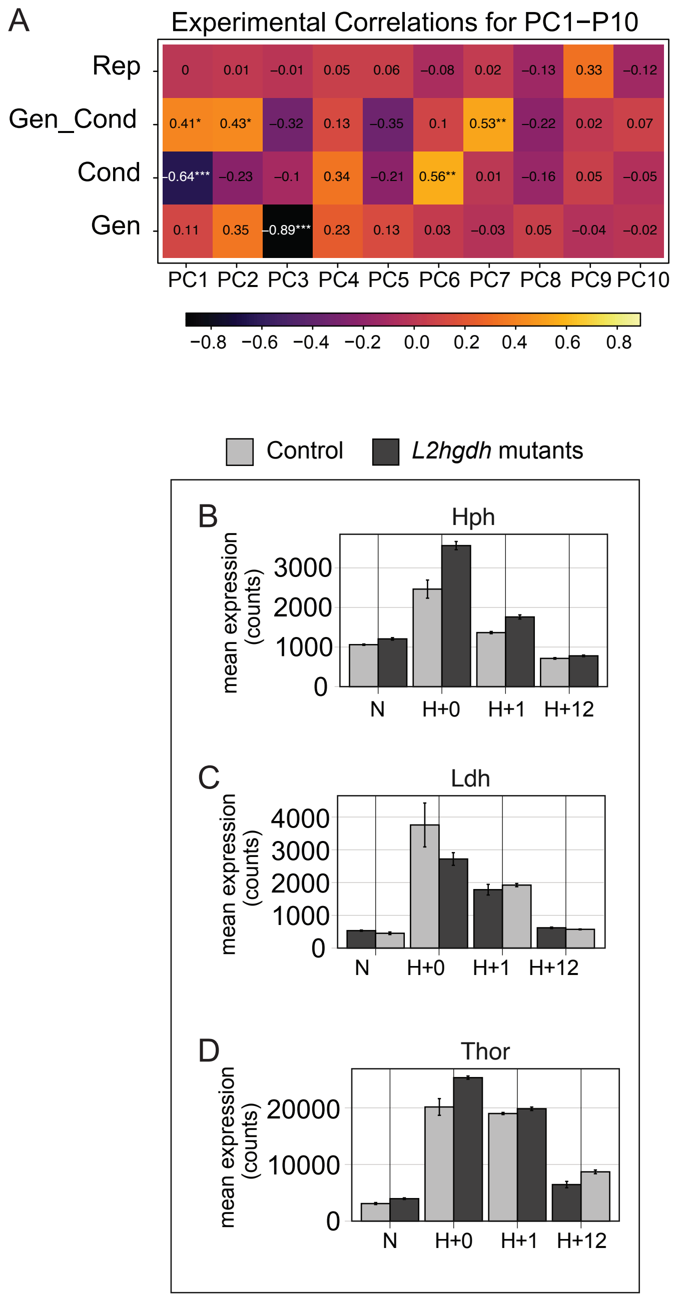


Supplemental Figure 11

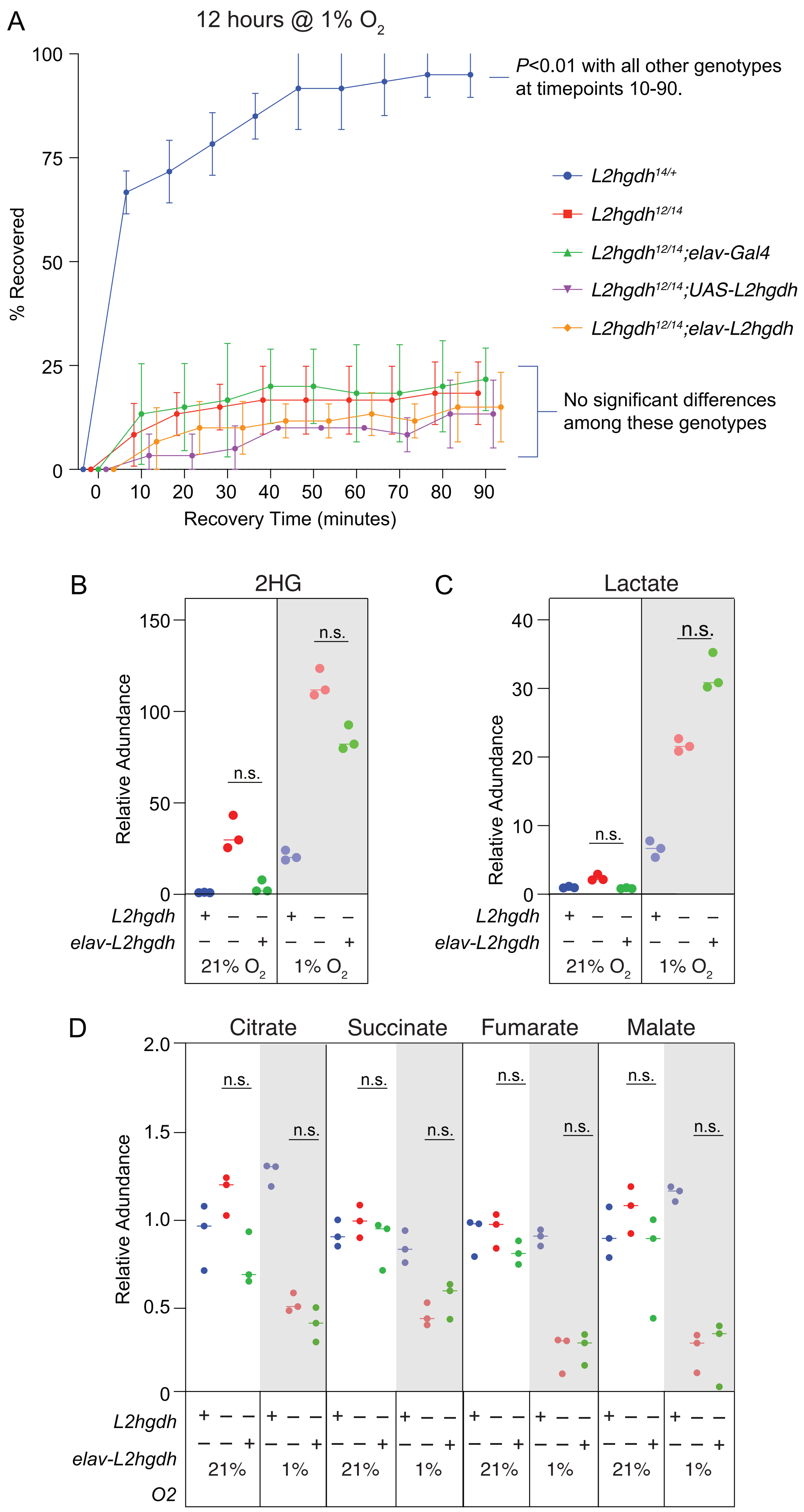


Supplemental Figure 12

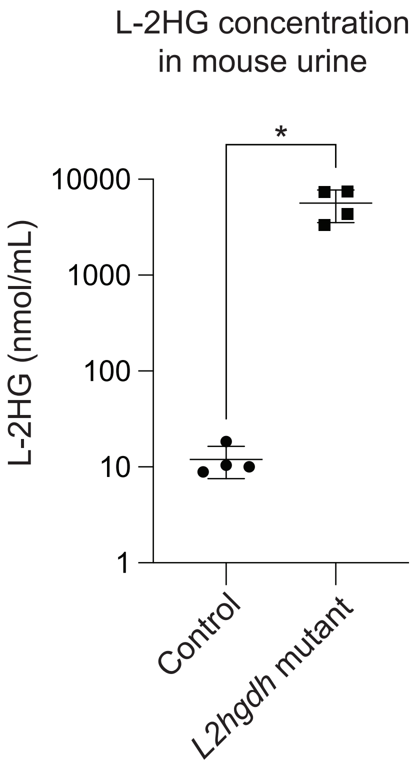


Supplemental Figure 13

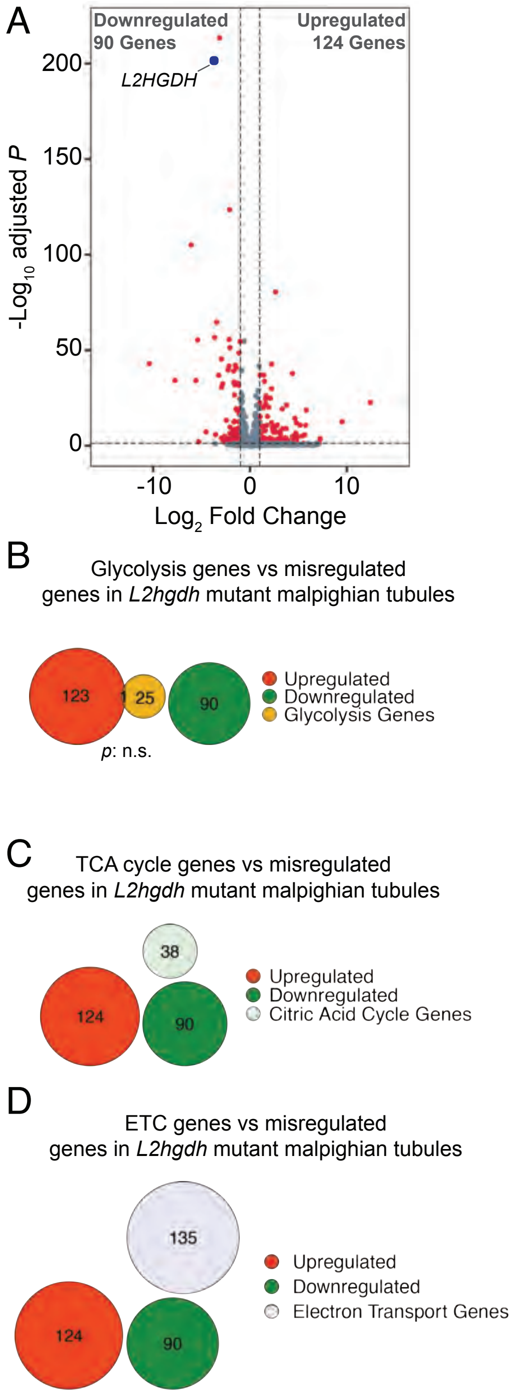
A
Downregulated
90 Genes
Upregulated
124 Genes
L2HGDH
200
150
100
50
0
-Log10 adjusted P
-10
0
10
Log2 Fold Change
B
Glycolysis genes vs misregulated
genes in L2hgdh mutant malpighian tubules
123
1
25
90
p: n.s.
Upregulated
Downregulated
Glycolysis Genes
C
TCA cycle genes vs misregulated
genes in L2hgdh mutant malpighian tubules
38
124
90
Upregulated
Downregulated
Citric Acid Cycle Genes
D
ETC genes vs misregulated
genes in L2hgdh mutant malpighian tubules
135
124
90
Upregulated
Downregulated
Electron Transport Genes

Figure S14
